# Calsyntenin-1 and calsyntenin-3 coordinate TGN exit of axonal cargoes

**DOI:** 10.64898/2026.06.15.732324

**Authors:** Derk Draper, Noortje Kersten, Shietela Sewnarain Sukul, Rebecca Jark, Inge Rothuis, Poorvi Ravindra, Aniruddha Mitra, Lukas C. Kapitein, Shawn Ferguson, Judith Klumperman, Ginny G. Farías

## Abstract

Neurons rely on precise biosynthetic protein transport from the soma to the axon and dendrites. Brain-enriched calsyntenins (CSTN1–3) are unique transmembrane adaptors that link cargo to kinesin-1 for transport. Yet, the neuronal distribution and interdependence of the three CSTN paralogs remain unclear. Here, we dissected their subcellular localization and contribution to biosynthetic protein transport. CSTN paralogs were predominantly localized to the TGN and axonal vesicles, with CSTN1 and CSTN3 showing higher expression levels than CSTN2. Depletion of either CSTN1 or CSTN3 affected axonal abundance of the other, suggesting that they function within the same pathway. Consistently, knockdown of CSTN1 or CSTN3, but not CSTN2, impaired the TGN exit and transport of multiple biosynthetic cargoes to the axon. Interestingly, most CSTN-positive vesicles exiting the TGN and in the axon were marked by the biosynthetic trafficking regulator RAB6A. Furthermore, we identified opposing, paralog-specific roles of CSTN1 and CSTN3 in regulating RAB6A levels at the TGN, thereby contributing to Golgi organization and axonal trafficking. Loss of CSTN1 reduced RAB6A and induced Golgi compaction, whereas loss of CSTN3 increased RAB6A and promoted Golgi dispersal. Together, we reveal that CSTN1 and CSTN3 have distinct, yet intersecting roles in the regulation of biosynthetic axonal transport.

## Introduction

Neuronal cells have a highly complex morphology, with distinct axonal and somatodendritic domains. To function properly, neurons need to establish and maintain distinct proteomes within each domain. Correct sorting of newly synthesized membrane proteins from the trans-Golgi network (TGN), located at the soma, to dendrites and the axon is crucial for this process (Bentley and Banker, 2016; Kersten and Farías, 2023). These biosynthetic proteins (further referred to as cargoes) are packaged into vesicles at the TGN and then selectively transported by microtubule-bound motors into the appropriate domain (Bentley and Banker, 2016). Direct motor-cargo binding is limited to a relatively small group of motor proteins, opposed to an enormous cargo diversity. Hence, adaptor proteins are required for motor-cargo recognition (Akhmanova and Hammer, 2010; Koppers and Farías, 2021).

Calsyntenins (CSTNs) are a family of adaptor proteins that are essential for neuronal development (Ponomareva et al., 2014; Um et al., 2014). Unlike conventional adaptor proteins, CSTNs have the unique feature that they are membrane proteins (Hintsch et al., 2002). This family comprises three proteins (CSTN1, CSTN2, CSTN3), which are almost exclusively expressed in brain tissue (Hintsch et al., 2002). Among this family, CSTN1 has been studied most extensively, whereas specific roles of CSTN2 and CSTN3 remain comparatively less well defined. CSTNs bind to the kinesin light chain-1 (KLC1) subunit of the axonal-directed motor kinesin-1 (KIF5A, KIF5B, and KIF5C), thereby regulating anterograde cargo transport to the axon (Konecna et al., 2006; Ludwig et al., 2009).

Besides their role in mediating axonal transport, CSTNs were originally described as dendritic postsynaptic proteins involved in synaptogenesis (Vogt et al., 2001; Hintsch et al., 2002; Um et al., 2014). They contain cadherin domains in their luminal / extracellular domain, which have been proposed to be part of trans-synaptic interactions with other adhesion molecules, like the axonally localized protein neurexin-1α (Lu et al., 2014). This dual function as both axonal trafficking adaptors and dendritic postsynaptic organizers raises intriguing questions about their primary site of action. Moreover, whether CSTN family members function independently or share common pathways remains unclear.

A well-characterized axonal cargo protein of CSTNs is the amyloid precursor protein (APP), a membrane protein involved in Alzheimer’s Disease (AD) pathogenesis (Ludwig et al., 2009; Steuble et al., 2012; Vagnoni et al., 2012). Aberrant localization of APP affects its proteolytic processing, thereby generating toxic Aβ oligomers and APP C-terminal fragments (APP-CTFs), which accumulate in AD patients (Yu et al., 2024; Luo et al., 2025). Consistent with the reported interaction with kinesin-1 (Konecna et al., 2006), CSTN1 predominantly mediates anterograde axonal trafficking of APP (Vagnoni et al., 2012). Importantly, postmortem AD brains show lower expression levels of CSTN1 (Vagnoni et al., 2012), while CSTN1 depletion increases APP levels at the TGN, suggesting a role of CSTN1 in APP exit from the TGN (Vagnoni et al., 2012). Although the three CSTN paralogs share strong structural similarities (Hintsch et al., 2002), it remains unclear whether CSTN2 and CSTN3 contribute equally to APP transport or whether this function is specific to CSTN1. More broadly, the extent to which additional cargo proteins depend on CSTNs for efficient axonal delivery remains largely unexplored.

In this study, we systematically examined the distribution and contribution of CSTN1-3 to biosynthetic protein trafficking in primary rat hippocampal neurons. Although CSTN2 was expressed at lower levels than its paralogs, all CSTNs were predominantly found in the TGN and axonal vesicles. We found that the loss of CSTN1 or CSTN3, but not CSTN2, impaired the axonal entry of RAB6A, a central regulator of post-Golgi transport (Storrie et al., 2012), alongside multiple biosynthetic cargoes. Notably, CSTN1 depletion decreased total RAB6A levels and induced Golgi compaction, whereas CSTN3 depletion increased somatic RAB6A levels and dispersed the Golgi stacks. Together, these data indicate that CSTN1 and CSTN3 exert opposite yet coordinated roles in the TGN exit of axonal cargoes, which could play an essential role in the development and maintenance of neuronal polarity.

## Results

### CSTNs localize to the Golgi region and in axonally-enriched vesicles

Since it remains unclear whether the CSTN paralogs localize to the dendritic and / or axonal domain (Hintsch et al., 2002; Konecna et al., 2006), we studied their endogenous subcellular localization in primary rat hippocampal neurons. For this purpose, we generated N-terminally tagged knock-in constructs using either the CRISPIE (CSTN1, CSTN3) (Zhong et al., 2021) or the ORANGE (CSTN2) (Willems et al., 2020; Droogers et al., 2022) systems (Fig. 1A-C; Fig. S1A) and investigated their distribution in neurons at *day-in-vitro* (DIV) 9 and 21 (Fig. 1D-F). Rat primary neurons at DIV9 are already polarized but lack mature synapses, while DIV21 neurons represent a more mature stage with fully developed synapses (Kaech and Banker, 2006).

**Fig. 1.**
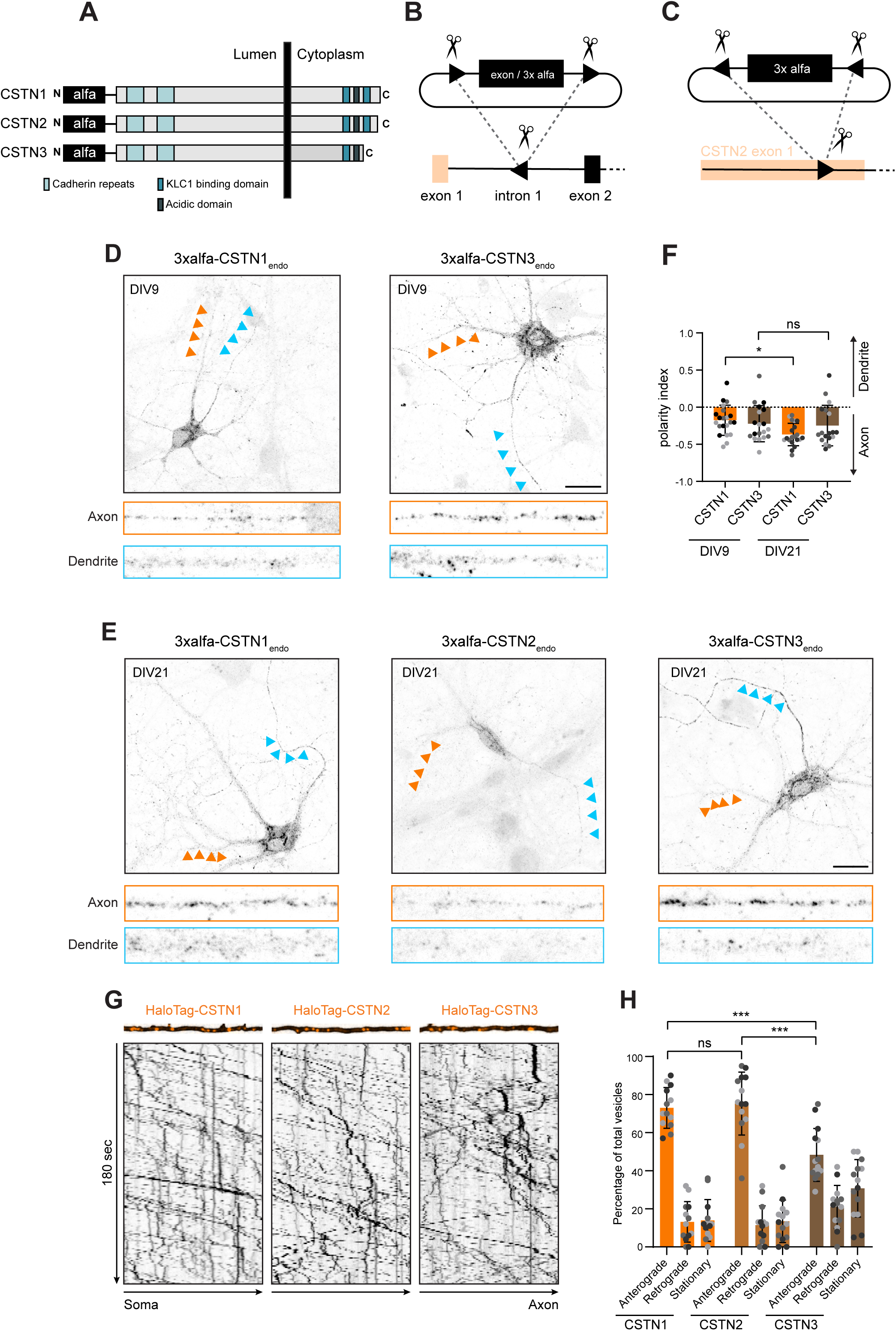
CSTNs predominantly localize to the Golgi region and axonal vesicles. (A) schematic presentation of CSTN1-3 including endogenous alfa tag. (B, C) Schematic presentation of knock-in strategy for CSTN1 and CSTN3, by CRISPIE (B) and CSTN2 by ORANGE (C). (D-E) Representative images of hippocampal neurons with endogenously-tagged CSTN1 or CSTN3 at DIV9 and endogenously tagged CSTN1-3 at DIV21. Cropped neurites are 30 µm. Scale bars: 10 µm. (F) Polarity index of CSTN signal at DIV9 and DIV21 for conditions shown in (D-E). (G) Representative still images and kymographs of live axons (30 µm) of DIV7 hippocampal neurons transfected on DIV4 with HaloTag-CSTN1, HaloTag-CSTN2 or HaloTag-CSTN3. Kymographs were obtained from 3-minute videos (1s/frame). (H) Directionality of CSTN1-3 compartments for conditions shown in (G). All data is presented as mean values ± SD. Dots in all dot plots indicate individual cells and replicates are indicated by different colors. In (F), we used the Kruskal-Wallis test followed by Dunn’s correction. In (H), we used two-way ANOVA followed by Sidak’s correction. *p value < 0.05, **p value < 0.01, ***p value < 0.001, ns = not significant.

At both developmental stages, we observed a robust immunofluorescence signal for endogenous CSTN1 and CSTN3 (Fig. 1D, E). Endogenous CSTN2 was not detectable at DIV9 and only slightly elevated at DIV21 (Fig. 1E). In the soma, all CSTN paralogs were enriched in the perinuclear region (Fig. 1D, E). Double-labelling of CSTN1 and CSTN3 with the cis-Golgi network (CGN) marker GM130 or trans-Golgi network (TGN) marker TGN38 revealed that they localized predominantly in the TGN (Fig. S1B, C), which is in agreement with previous reports (Ludwig et al., 2009; Vagnoni et al., 2012).

Next, we studied the distribution of the CSTNs between the dendritic and axonal domains. We calculated the polarity index for CSTN1 and CSTN3 on DIV9 and DIV21 ((PI = [intensity dendrite − intensity axon] / [intensity dendrite + intensity axon]), in which PI = 0, unpolarized; PI >0, dendritic; PI <0, axonal) (Kapitein et al., 2010). Interestingly, both CSTN1 and CSTN3 were highly polarized to the axonal domain at both DIV9 and DIV21 (Fig. 1F). CSTN2 displayed numerous axonal puncta, but a reliable quantification was prevented by the poor signal (Fig. 1E).

Since CSTN1 and CSTN2 contain two kinesin-1 binding sequences (KBS) whereas CSTN3 only contains one (Fig. 1A) (Konecna et al., 2006), we wondered whether their dynamics along the axon differ. To assess this, we performed live-cell imaging in neurons overexpressing individual CSTN paralogs with an N-terminal HaloTag (Fig. 1G). We acquired 3-minute videos for each, and generated kymographs to study their directionality of movement along the axon. We observed that all the paralogs preferentially moved in the anterograde direction (Fig. 1G, H; Video S1). CSTN3, however, displayed less frequent anterograde movement compared to CSTN1 and CSTN2 (Fig. 1G, H; Video S1), likely due to the presence of only one KBS.

Together, these results show that CSTN paralogs localize to the TGN and are present in axonal vesicles that undergo preferential anterograde transport.

### CSTN paralogs colocalize in axonal vesicles and are interdependent for their axonal localization

Given the preferential axonal distribution of all CSTN paralogs, we wondered whether they are present in the same or distinct axonal vesicle populations. To address this, we examined their colocalization by expressing the same CSTN constructs as above, with either an mNeonGreen or HaloTag. As a positive control, simultaneous expression of two CSTN1 constructs with different tags was included, which resulted in a high level of overlap (Fig. 2A). Likewise, both CSTN2 and CSTN3 showed a high degree of colocalization with CSTN1 (Fig. 2A, B). Importantly, the number of CSTN-positive vesicles was not changed upon overexpression compared to endogenously labeled CSTNs (Fig. S2A). We conclude from these data that the three CSTN paralogs colocalize in the same axonal vesicles.

**Fig. 2.**
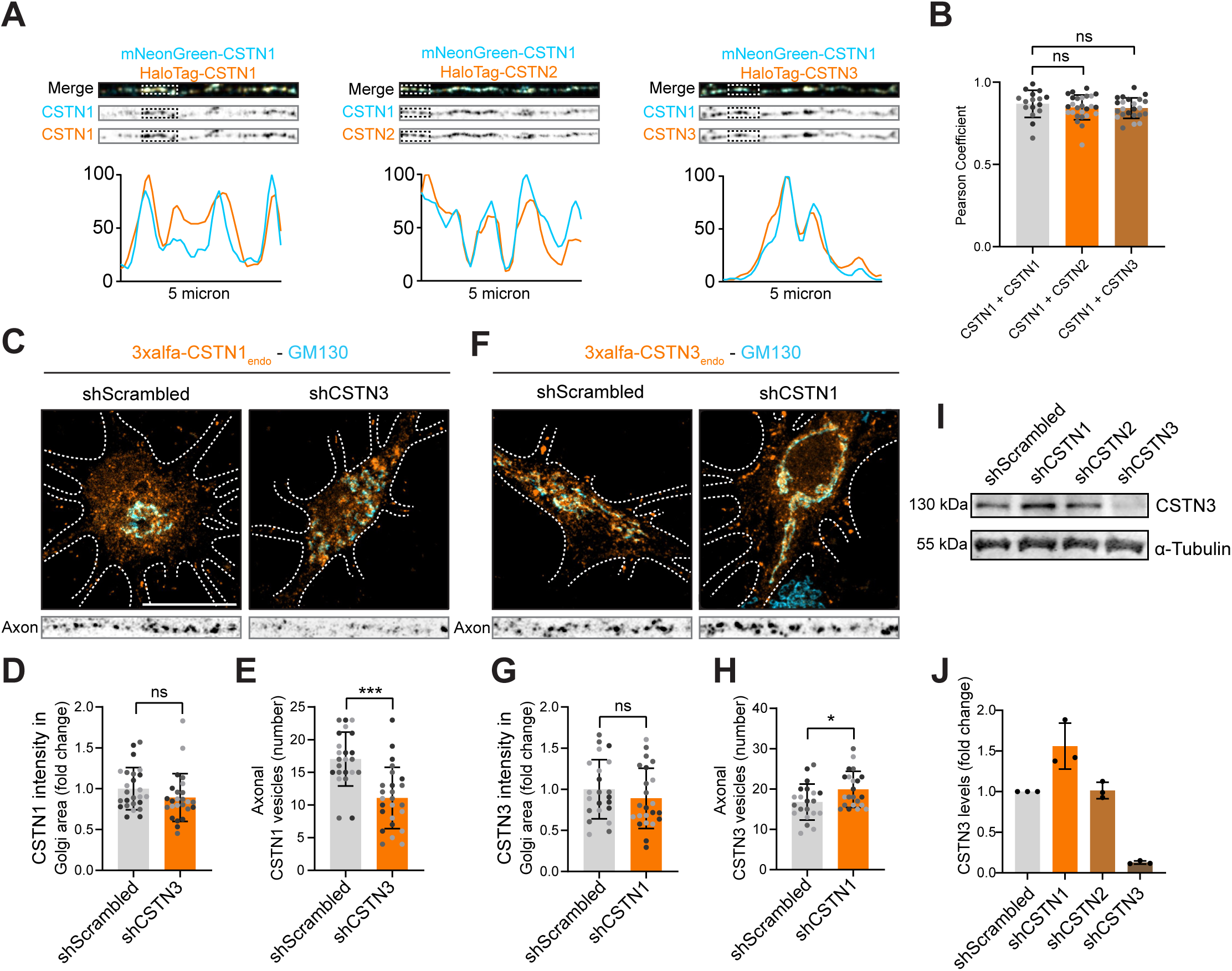
CSTN paralogs colocalize in axonal vesicles and are interdependent for their axonal localization. (A) Representative images of axons (30 µm) of DIV8 hippocampal neurons transfected on DIV7 with mNeonGreen-CSTN1 and / or HaloTag-CSTN1 and / or HaloTag-CSTN2 and / or HaloTag-CSTN3 (indicated in the figure). Profile plots of indicated square boxes show the intensity as a fraction of the maximum intensity of that channel. (B) Pearson coefficient for conditions shown in (A). (C) Representative images of DIV9 hippocampal neurons with endogenously tagged CSTN1 or CSTN3 (orange), transduced at DIV4 with lentivirus containing shScrambled-GFP, shCSTN1-GFP or shCSTN3-GFP, and stained for GM130 (blue). Axon crops (30 µm) are indicated below. Scale bar: 20 µm. CSTN intensity in Golgi region (D, G) and number of axonal CSTN compartment (per 30 µm) (E, H) are indicated in graphs below. All data is presented as mean values ± SD. Dots in all dot plots indicate individual cells and replicates are indicated by different colors. In (B), Kruskal-Wallis test was used followed by Dunn’s correction. In (D), Mann-Whitney test was used. In (E, G, H), unpaired t test was used. *p value < 0.05, **p value < 0.01, ***p value < 0.001, ns = not significant.

Because of the high degree of colocalization between the paralogs, we asked whether their functions might be interdependent. To address this, we generated lentiviral shRNA constructs targeting individual CSTNs. After validating the efficacy of CSTN depletion by qPCR (Fig. S2B-D), we examined the distribution of endogenous CSTN1 and CSTN3 following depletion of the other, i.e. CSTN3 and CSTN1, respectively. The low expression levels of CSTN2 prevented us from doing similar assays on this paralog. For both CSTN1 and CSTN3, no changes were observed in their localization to the Golgi area upon depletion of the other paralog (Fig. 2C, D, F, G). Intriguingly, CSTN3 depletion reduced the number of axonal CSTN1-positive vesicles, whereas CSTN1 depletion led to an increase in axonal CSTN3-positive vesicles (Fig. 2C, E, F, H). We next performed Western blot analysis for endogenous CSTN3 on lysates obtained from cortical neurons transduced with shScramble or shCSTN constructs (Fig. 2I). Consistent with the increase in axonal CSTN3, we observed elevated CSTN3 expression levels in CSTN1-depleted neurons (Fig. 2I-J). In addition, CSTN2 depletion had no effect on total CSTN3 levels (Fig. 2I-J), consistent with their lower expression levels (Fig. 1E). We did not perform immunoblotting of endogenous CSTN1, because of the lack of a suitable antibody.

Altogether, we show that all CSTN paralogs colocalize in the same axonal vesicles. Upon CSTN1 depletion CSTN3 levels in the axon are upregulated, while vice versa CSTN3 depletion decreases axonal CSTN1 levels. Together, these data suggest that CSTN1 and CSTN3 have complementary functions within the same axonal trafficking pathway.

### CSTN1 and CSTN3 mediate Golgi-to-axon transport of APP

Previous studies have shown that CSTN1 regulates axonal transport of APP (Steuble et al., 2012; Vagnoni et al., 2012). Since we observe that CSTN paralogs highly colocalize in the same axonal vesicles and that their distribution depends on the presence of each other, we wondered whether CSTN2 and CSTN3 are also involved in APP trafficking to the axon. To this end we co-expressed N-terminally tagged APP and the CSTN paralogs (Fig. 3A, Video S2). This clearly showed the presence of APP in CSTN-positive axonal vesicles. Interestingly, the vast majority of CSTN-positive / APP-positive vesicles exhibited anterograde transport along the axon (Fig. 3A-B), consistent with our data on CSTN preferential anterograde movement (Fig. 1; Fig. 2).

**Fig. 3.**
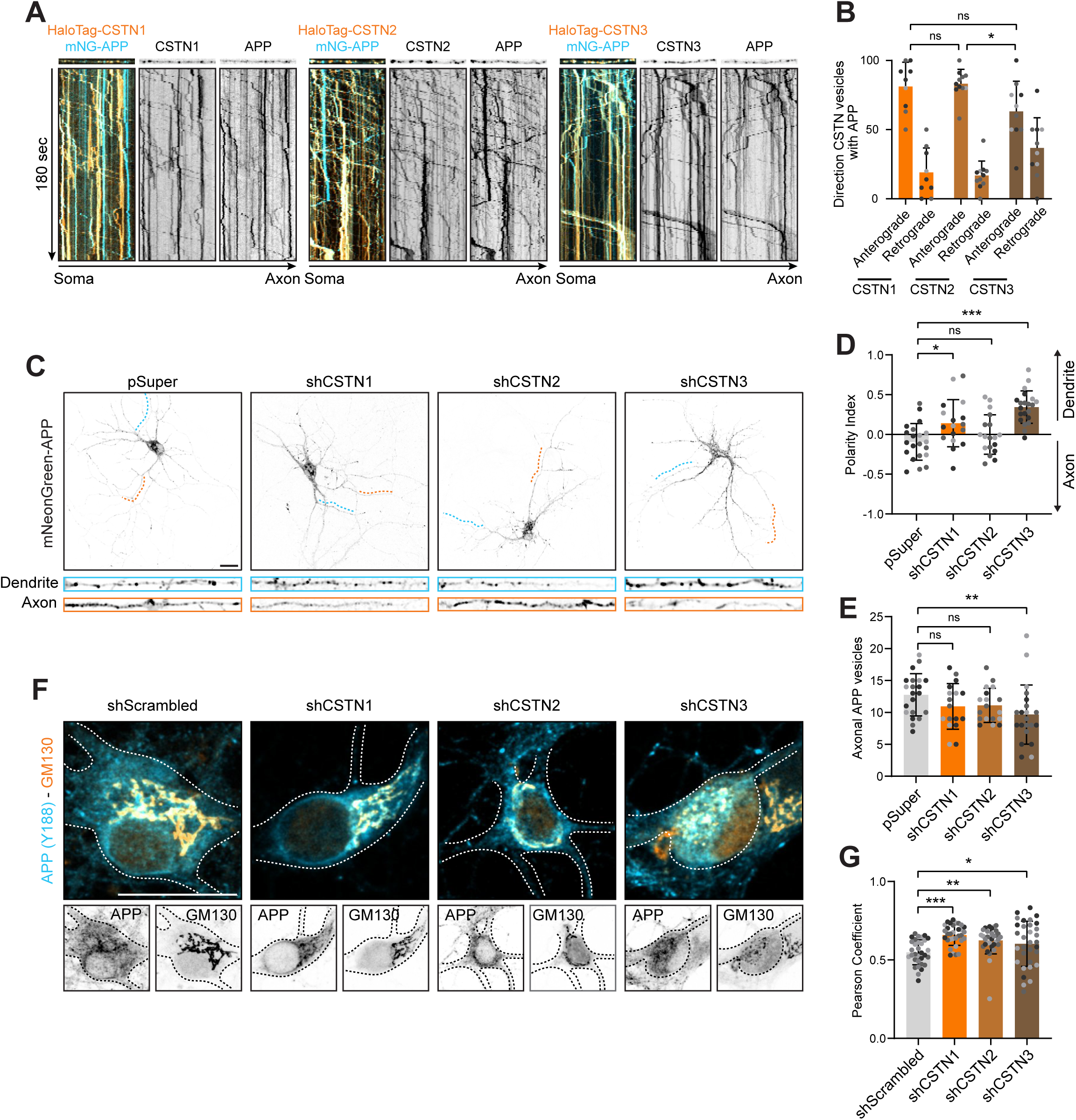
CSTN1 and CSTN3 are required for Golgi-to-axon transport of APP. (A) Representative still images and kymographs of live axons (30 µm) of DIV7 hippocampal neurons transfected on DIV4 with HaloTag-CSTN1, HaloTag-CSTN2 or HaloTag-CSTN3 (orange) together with mNeonGreen-APP (blue). Kymographs were obtained from 3-minute videos (1s/frame). (B) Directionality of colocalized APP and CSTN1-3 compartments for conditions shown in (A). (C) Representative images of DIV8 hippocampal neurons transfected on DIV4 with mNeonGreen-APP and empty pSuper or pSuper-shRNA for CSTN1-3. Scale bar: 20 µm. Cropped dendrites and axons indicate a region of 30 µm. Polarity index (D) and number of axonal vesicles (per 30 µm) (E) for conditions shown in (C). (F) Representative images of DIV8 hippocampal neurons transfected with shScrambled-GFP or shCSTN1-3-GFP and labelled for APP (blue) and GM130 (orange). Scale bar: 20 µm. Individual channels are indicated below. All data is presented as mean values ± SD. Dots in all dot plots indicate individual cells and replicates are indicated by different colors. In (B), two-way ANOVA was used followed by Sidak’s correction. In (D), one-way ANOVA was used followed by Dunnett’s correction. In (E,G), Kruskal-Wallis test was used followed by Dunn’s correction. *p value < 0.05, **p value < 0.01, ***p value < 0.001, ns = not significant.

Then, we assessed the requirement of the different paralogs for axonal APP trafficking. For this, we expressed tagged APP in CSTN-depleted neurons and quantified the polarity index between axon and dendrites. Consistent with previous reports (Allinquant et al., 1994; DeBoer et al., 2014; Draper et al., 2026), APP distribution was slightly polarized to the axonal domain in control neurons (Fig. 3C-D). In contrast, depletion of either CSTN1 or CSTN3 reduced axonal APP signal, resulting in a shift of APP polarity towards dendrites (Fig. 3C-D). This effect was more pronounced in CSTN3 than in CSTN1-depleted neurons and APP distribution was not altered in CSTN2-depleted cells (Fig. 3C-D). We also quantified the number of axonal APP-positive vesicles and observed a significant reduction of axonal APP vesicles in CSTN3-depleted cells and a trend of a reduction in CSTN1-depleted cells (Fig. 2E).

Previously, CSTN1-positive / APP-positive tubules have been observed in the Golgi region of non-polarized COS7 cells (Ludwig et al., 2009). Also, CSTN1 depletion has been shown to increase APP signal at the Golgi region in neurons, suggesting impaired TGN exit (Vagnoni et al., 2012). However, it remains unclear whether the other CSTN paralogs also influence the Golgi distribution of APP in neurons. To address this, we knocked down CSTNs in neurons and labeled for endogenous APP and the CGN marker GM130 (Fig. 3F) and measured the colocalization between APP and GM130 (Fig. 3G). In control neurons, APP showed moderate Golgi localization (Fig. 3F, G), consistent with previous data (Tan and Gleeson, 2019; Draper et al., 2026). Interestingly, depletion of all three CSTN paralogs consistently increased the colocalization of APP and GM130 in the Golgi region (Fig. 3F, G).

Together, these data show that all CSTN paralogs co-distributed with axonal anterograde APP vesicles. Depletion of CSTN1 and CSTN3 resulted in decreased axonal APP and a shift in polarity towards dendritic APP, while depletion of all paralogs increased Golgi-localized APP. These results suggest a role of CSTN paralogs in the efficient exit of biosynthetic APP from the TGN and for its transport into the axon.

### CSTN1 and CSTN3 mediate biosynthetic axonal cargo trafficking

To further study the role of CSTNs in the biosynthetic route of APP, we applied the retention using selective hooks (RUSH) tool (Fig. 4A) (Boncompain et al., 2012). In short, we coupled APP to a streptavidin binding protein (SBP) together with mNeonGreen to allow its visualization. We co-expressed this construct with the ER-specific strep-KDEL hook, which leads to ER retention of newly synthesized APP. Upon addition of biotin, APP is released from the ER, allowing visualization of its itinerary in a synchronized manner (Draper et al., 2026). For simplicity, we will generally refer to this module as RUSH-POI (POI: protein of interest). First, we co-expressed RUSH-APP together with the individual CSTNs and investigated their co-trafficking from the TGN (Fig. 4B; Video S3). For this purpose, we first used HeLa cells, as their flat morphology allows for better visualization of budding events from the TGN. For all three paralogs, we observed multiple co-budding events of CSTN and RUSH-APP (Fig. 4B; Video S3). Also in neurons, RUSH-APP was efficiently retained in the ER in the absence of biotin (Fig. S3A; Video S4). After biotin addition, RUSH-APP was sorted from the TGN directly to the axon 30 minutes post release (Fig. S3A; Video S4) and these vesicles highly co-localized with all CSTN paralogs along the axon (Fig. 4C). The synchronized released by the RUSH system leads to high local levels in the Golgi system. To exclude that our observations were caused by overexpression, we restricted the amount of RUSH-APP exiting the ER by adding NeutrAvidin 5 min post-biotin addition. NeutrAvidin competes with biotin for binding to SBP (Fig. S3B). This short biotin pulse resulted in a similar high colocalization between CSTN3 and biosynthetic APP as in neurons exposed to biotin for 1h (Fig. S3B, C), suggesting this high degree of colocalization is not caused by Golgi overloading.

**Fig. 4.**
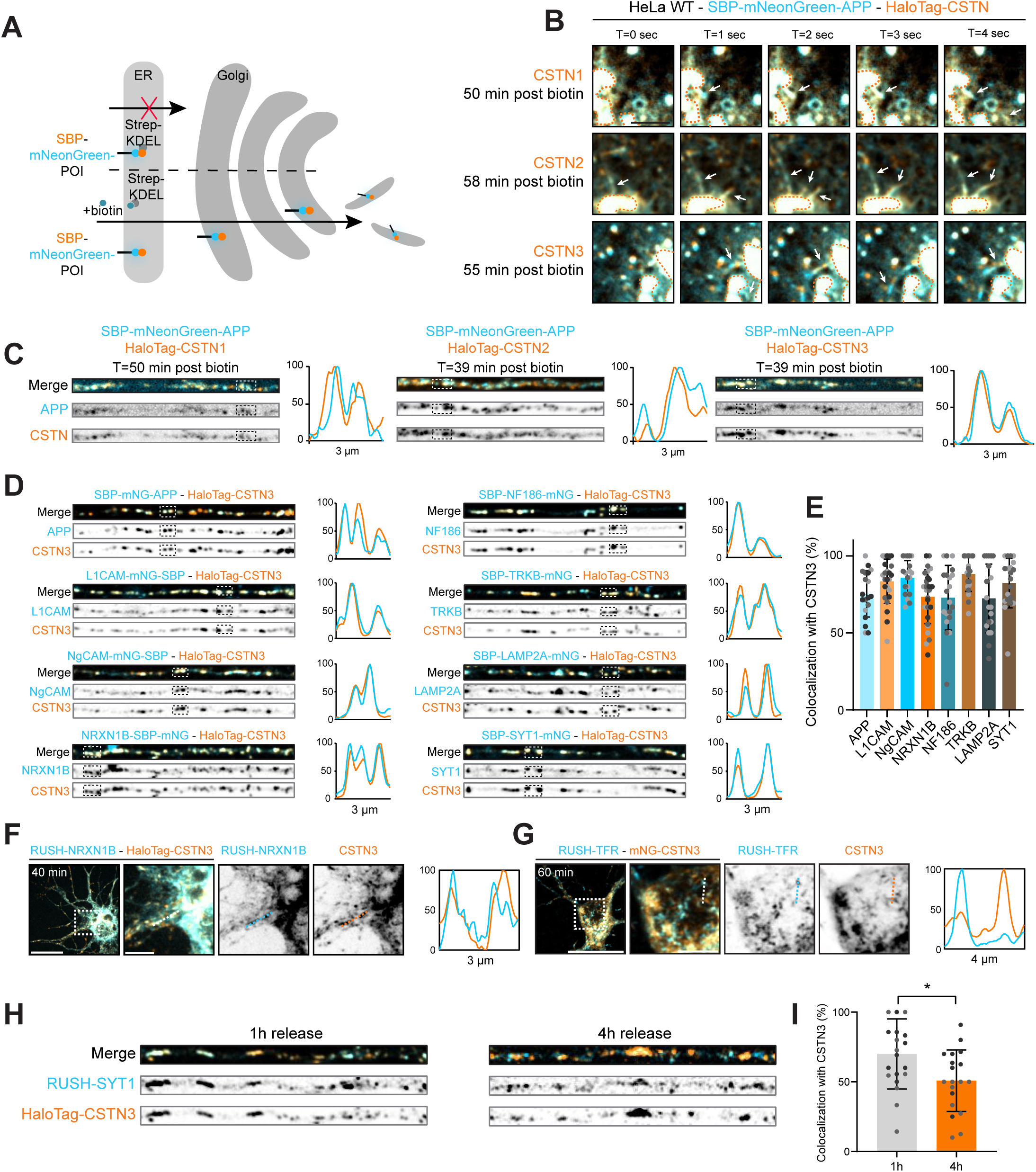
CSTN1-3 colocalize with multiple biosynthetic axonal cargoes. (A) Schematic of the RUSH system for retention and release of protein of interest (POI) from the ER. (B) Representative stills derived from live HeLa cells co-transfected with RUSH-APP and HaloTag-CSTN1-3 constructs. Orange dotted lines indicate the Golgi region and white arrows indicate individual budding events. Scale bar: 5 µm (C) Representative stills derived from axons (30 µm) of live DIV8 hippocampal neurons co-transfected on DIV7 with RUSH-APP and HaloTag-CSTN1-3 constructs. Intensity profiles of boxed regions are indicated on the right beside respective stills. (D) Representative crops of axons (30 µm) from DIV8 hippocampal neurons co-transfected on DIV7 with HaloTag-CSTN3 and RUSH-POI constructs. RUSH-POI was released for 1h Intensity profiles of boxed regions are indicated on the right beside respective crops. (E) Quantification of percentage of RUSH-POI compartments containing HaloTag-CSTN3 from conditions in (D). (F, G) Representative images derived from DIV8 neurons co-transfected on DIV7 with RUSH-NRXN1B (F) or RUSH-TFR (G) and CSTN3 constructs. Scale bars: 20 µm (overview), 5 µm (zoom). Magnification of boxed region (right). Intensity profiles of lines are indicated on the right beside respective images. (H) Representative crops of axons (30 µm) from DIV8 hippocampal neurons co-transfected on DIV7 with HaloTag-CSTN3 and RUSH-SYT1 construct. RUSH-SYT1 was either released for 1h (left) or 4h (right). (I) Quantification of percentage of RUSH-SYT1 compartments containing HaloTag-CSTN3 from conditions in (H). All data is presented as mean values ± SD. Dots in all dot plots indicate individual cells and replicates are indicated by different colors. Times indicate time post biotin addition in all images. In (I), unpaired t-test was used. *p value < 0.05, **p value < 0.01, ***p value < 0.001, ns = not significant.

We then wondered whether CSTNs might also play a role in the biosynthetic transport of other cargoes into the axon. To this end, we studied the trafficking of biosynthetic adhesion molecules (L1CAM, NgCAM, NRXN1β, NF186), the growth receptor TRKB, the lysosomal protein LAMP2A, and the synaptic protein SYT1. We co-expressed RUSH constructs of these proteins together with CSTN3 (Fig. 4D, E), because of its high expression and dominant function in the previous assays. At 1h post-release from the ER, we observed high colocalization (70-90%) between all studied biosynthetic cargoes and CSTN3 in the axon (Fig. 4D, E). In addition, live-cell imaging of RUSH-LAMP2A and RUSH-NRXN1β revealed a high degree of co-transport with CSTN3 in both soma and anterogradely along the axon (Fig. 4F; Fig. S3D, E). Importantly, we observed little colocalization between the somatodendritic cargo TFR and CSTN3 (Fig. 4G) (Farías et al., 2012, 2015), indicating specificity of CSTN3 for axonal cargo. Notably, colocalization of RUSH-SYT1 with CSTN3 reduced after 4h compared to 1h release, indicating a specific role of CSTNs mainly in biosynthetic vesicular transport, as RUSH-SYT1 reaches synaptic vesicles at 4h (Fig. 4H, I) (Li et al., 2024; Nguyen et al., 2025).

To further investigate the role of CSTN paralogs in transport from the TGN into the axon, we depleted each paralog individually and measured cargo intensity in the perinuclear region and the proximal axon 1h post-release (Fig. 5A). Upon knockdown of CSTN1 as well as of CSTN3, biosynthetic APP, L1CAM, TRKB and LAMP2A signals were increased in the Golgi area (Fig. 5B-I). This was paired with signal reduction in the axon (Fig. 5B-I), implying impaired biosynthetic trafficking from the Golgi to the axon. This impaired trafficking was even more pronounced in CSTN3-depleted cells (Fig. 5B-I). CSTN2 depletion did not result in a Golgi block of these cargo proteins, and showed normal levels in the axon, which may be explained by its lower expression levels (Fig. 5B-I; Fig 1E). These results were validated by immuno-labelling of fixed, RUSH-L1CAM expressing neurons 1h post-release, with the CGN marker GM130 and axonal marker TRIM46 (Fig. S3F). This confirmed reduced axonal localization of biosynthetic L1CAM and accumulation at the Golgi in shCSTN1 and shCSTN3 expressing neurons.

**Fig. 5.**
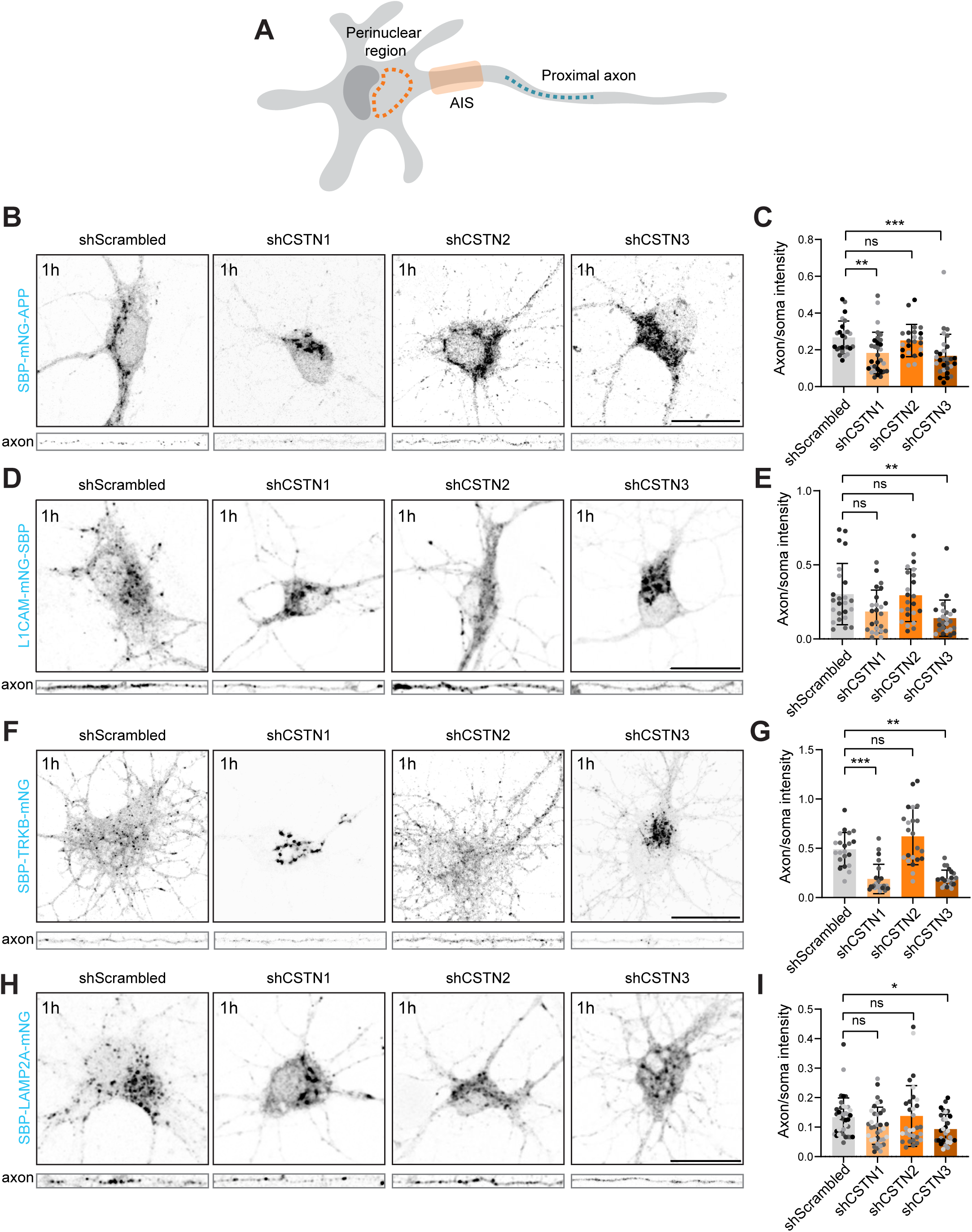
CSTN1 and CSTN3 are required for biosynthetic axonal cargo trafficking. (A) Schematic depicting regions quantified in B-I. (B, D, F, H) Representative images of soma and axon (50 µm) of DIV7-8 hippocampal neurons transfected at DIV4 with RUSH-mNeonGreen-APP (B), L1CAM-mNeonGreen-RUSH (D), RUSH-TRKB-mNeonGreen (F), RUSH-LAMP2A-mNeonGreen (H) and shScrambled, shCSTN1, shCSTN2 or shCSTN3. RUSH-POI was released in all experiments for 1h. Scale bars: 20 µm. (C, E, G, I) Ratio of the mean intensity in the axon divided by the mean intensity in the perinuclear area. All data is presented as mean values ± SD. Dots in all dot plots indicate individual cells and replicates are indicated by different colors. Times indicate time post biotin addition in all images. In (C, E, G, I), Kruskal-Wallis test was used followed by Dunn’s correction for multiple comparisons. *p value < 0.05, **p value < 0.01, ***p value < 0.001, ns = not significant.

Together, we found that CSTN1 and CSTN3 are involved in TGN exit of different biosynthetic cargoes and their subsequent, polarized transport into the axon.

### CSTN1 and CSTN3 coordinate the transport of RAB6A-positive vesicles to the axon by regulating RAB6A levels at the soma

Since multiple axonal cargoes depend on CSTNs for TGN exit and transport into the axon, we wondered whether CSTNs were involved in a RAB6A-dependent vesicular transport pathway to the axon. RAB6A is a GTPase with a general role in the formation TGN carriers and post-TGN transport of a broad range of biosynthetic proteins (Grigoriev et al., 2007; Storrie et al., 2012; Zahavi et al., 2021). By immuno-fluorescence we stained for endogenous RAB6A and quantified its colocalization with endogenously expressed CSTN1 and CSTN3, focusing on the axon (Fig. 6A, B). This showed that most axonal RAB6A puncta were positive for each paralog and vice versa, with greatly overlapping localization patterns (Fig. 6A, B).

**Fig. 6.**
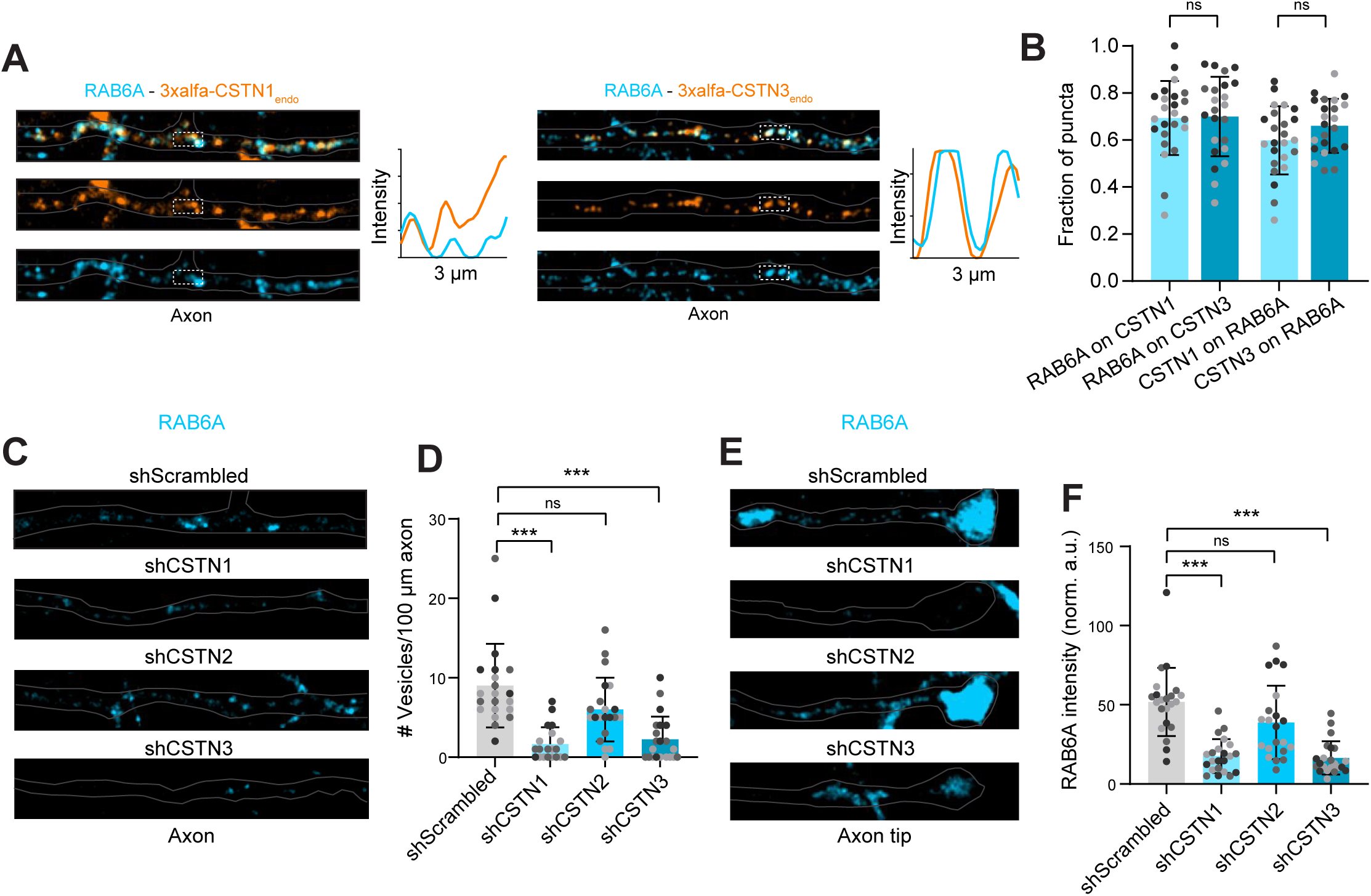
CSTN1 and CSTN3 associate with and mediate axonal transport of RAB6A-positive vesicles. (A) Representative images of axons (30 µm) of neurons with endogenously tagged CSTN1 or CSTN3 (orange), stained for RAB6A (blue). Profile plots of indicated boxed regions show intensity in arbitrary units. (B) Fraction of CSTN puncta positive for RAB6A and vice versa in 50 µm of axon. (C, E) Representative images of axons (30 µm) and axon tips (20 µm) of DIV7 neurons transfected at DIV3 with shScrambled-GFP, shCSTN1-GFP, shCSTN2-GFP or shCSTN3-GFP, immunolabeled for RAB6A. (D) Number of vesicles along 100 µM of axon. Statistical analysis by Kruskal-Wallis test. (F) RAB6A intensity (normalized arbitrary units) measured in the distal region of the axon tip. All data is presented as mean values ± SD. Dots in all dot plots indicate individual cells and replicates are indicated by different colors. In (B), one way ANOVA with Sidak’s correction was used. In (D, F), Kruskall-Wallis followed by Dunn’s correction was used. In *p value < 0.05, **p value < 0.01, ***p value < 0.001, ns = not significant.

Next, we knocked down the three CSTN paralogs and investigated the effect on the number of RAB6A vesicles along the axon (Fig. 6C-F). In control neurons, the number of axonal RAB6A puncta was strikingly higher than in dendrites, whereby RAB6A accumulated in the axonal tips, emphasizing a predominant role in axonal trafficking (Fig. 6E, F; Fig. S4A, B). Consistent with our RUSH experiments (Fig. 5), knockdown of either CSTN1 or CSTN3 decreased the number of RAB6A vesicles along the axon and in axon tips, while depletion of CSTN2 had no effect (Fig. 6C-F). Depletion of CSTN1 using an additional shRNA, targeting a different sequence, produced a similar reduction of RAB6A puncta (Fig. S4C, D).

In the soma, RAB6A colocalized with both CSTN1 and CSTN3 in the Golgi region and some nearby vesicles (Fig. 7A), suggesting that their association is initiated at the level of the TGN. Intriguingly, expression of both shRNAs targeting CSTN1 caused a pronounced reduction of RAB6A labeling in the perinuclear region (Fig. 7B, C; Fig. S5A, B). This decrease in fluorescent RAB6A signal was explained by a reduction in total RAB6A expression levels, as found by Western blot analysis (Fig. S5C). Surprisingly, depletion of CSTN3 showed a distinct phenotype, leading to an increased RAB6A signal in the soma, without any effect on total RAB6A expression levels (Fig. 7B, C; Fig. S5C). Of note, the RAB6A signal presented in both CSTN1 and CSTN3 depleted neurons was mainly associated with the Golgi region, with no apparent distribution to somatic or axonal vesicles (Fig. 7B). Overexpression of an shRNA-resistant HaloTag-CSTN3 construct in CSTN3-depleted neurons restored RAB6A levels in the soma to normal (Fig. S5D, E). Unexpectedly, however, RAB6A levels were also rescued by expressing a CSTN3 variant carrying a mutation that disrupts binding to KLC1 (Fig. S5D, E) (Konecna et al., 2006). These results suggest that the increase of somatic RAB6A upon CSTN3 knockdown is independent of its role in kinesin-1 binding. Of note, in contrast to RAB6A, GM130 levels were reduced in CSTN3 knockdown neurons compared to control neurons (Fig. 7B, D), which may potentially point to altered Golgi positioning or organization (Puthenveedu et al., 2006).

**Fig. 7.**
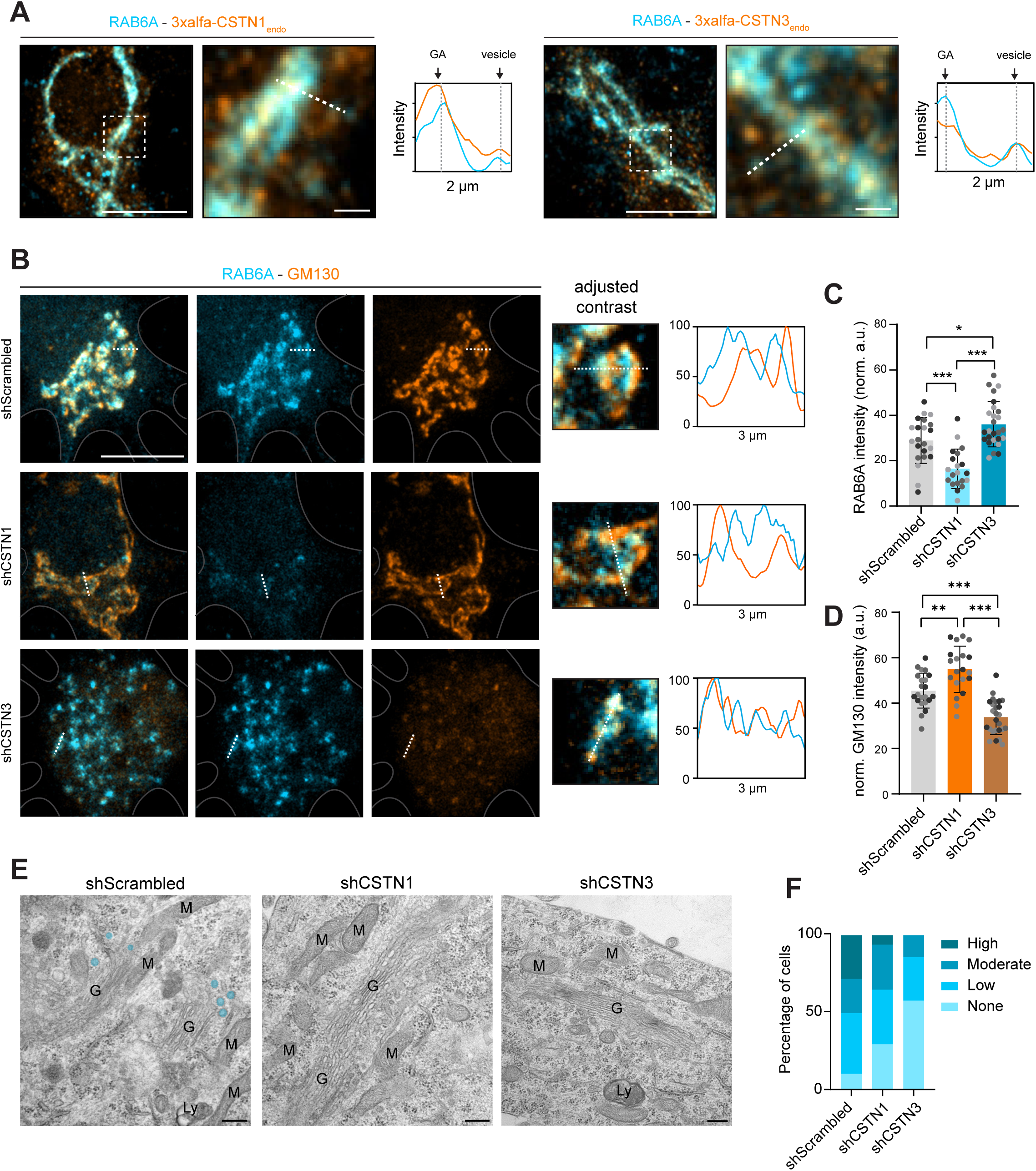
CSTN1 and CSTN3 conversely regulate RAB6A levels in the soma. (A) Representative images of DIV9 neurons with endogenously tagged CSTN1 or CSTN3, stained for RAB6A. Profile plot shows the intensity in arbitrary units. Scale bar of overview: 10 µm, scale bar of zoom: 1 µm. (B) Representative images of the soma of DIV7 neurons transfected at DIV3 with shScrambled-GFP, shCSTN1-GFP or shCSTN3-GFP, stained for RAB6A (blue) and GM130 (orange). Scale bar: 10 µm. Profile plots show the intensity as a percentage of the maximum value per channel. (C, D) RAB6A (C) and GM130 (D) intensity measured in the perinuclear region (normalized arbitrary units). (E) Representative EM images of DIV8 hippocampal neurons transduced at DIV4 with lentivirus with shScrambled-GFP, shCSTN1-GFP or shCSTN3-GFP. (F) Percentage of cells containing either none, low, moderate or high vesiculation at the TGN side of the Golgi complex. Detailed description can be found in the methodology section and visual description of categories in Fig. S6F. Scale bar: 200 nm. Data is presented as mean values ± SD in (C, D). Dots in all dot plots indicate individual cells and replicates are indicated by different colors in (C, D). In (C, D), one-way ANOVA with Tukey’s correction was used. *p value < 0.05, **p value < 0.01, ***p value < 0.001, ns = not significant.

To examine whether the effects on RAB6A, caused by CSTN1 and CSTN3 knockdowns, led to impaired vesicle budding from the TGN we next performed electron microscopy (EM). We quantitated the number of vesicular-tubular membrane profiles in the TGN area, including non- and clathrin-coated vesicles but excluding typical endosomes and COP-vesicles (Fig. S5F). We observed a reduction in the vesicles surrounding the TGN upon depletion of either CSTN1 or CSTN3 (Fig. 7E-F), consistent with an impaired biogenesis of vesicles at the TGN (Fig. 5A-I).

To study whether RAB6A presence at the TGN could be directly regulated by CSTN1 and CSTN3, we investigated their binding to RAB6A. We expressed the cytosolic domains of CSTN1 and CSTN3 coupled to GFP in HEK293T cells and exposed derived cell lysates to purified and immobilized GST or RAB6A-GST (Fig. S5G, H). As positive control, we evaluated RAB6A-GST binding to the dynein associated protein Bicaudal D cargo adaptor 2 (BICD2) (Fig. S5I), a well characterized interactor of RAB6A (Matanis et al., 2002). We indeed observed BICD2 and RAB6A interaction, but CSTN1 and CSTN3 did not directly interact with RAB6A (Fig. S5H-I).

Together, these results indicate that CSTN1 and CSTN3 both have a major yet functionally different role in the RAB6A-dependent transport pathway into the axon. Our data show that CSTN1 depletion decreases and CSTN3 depletion increases RAB6A levels in the soma, both leading to impaired vesicle formation at the TGN.

### CSTN1 and CSTN3 regulate Golgi positioning through RAB6A

Altered RAB6A levels in the soma of CSTN1 or CSTN3 depleted cells might have consequences for Golgi positioning, since RAB6A has been shown to play a role in Golgi organization in non-neuronal cell lines (Sun et al., 2007; Storrie et al., 2012). For this reason, we studied the role of CSTN paralogs on Golgi morphology. We first assessed the distribution of the CGN marker GM130 in control and CSTN-depleted neurons (Fig. 8A). In control cells, GM130 was predominantly localized to the perinuclear region, appearing as a network of punctae and tubules (Fig. 8A, B). In CSTN1-depleted neurons, the number of GM130-positive puncta was reduced, and the staining appeared condensed into elongated Golgi cisternae (Fig. 8A, B). In contrast, depletion of CSTN3 led to an increased number of GM130 puncta (Fig. 8A, B). Expansion microscopy further revealed the presence of numerous small GM130-positive structures upon CSTN3 depletion, whereas CSTN1 depletion led to a compacted tubular network (Fig. S6A). Despite these alterations in organization, the total GM130-positive area per cell remained unchanged upon depletion of either CSTN1 or CSTN3 (Fig. S6B).

**Fig. 8.**
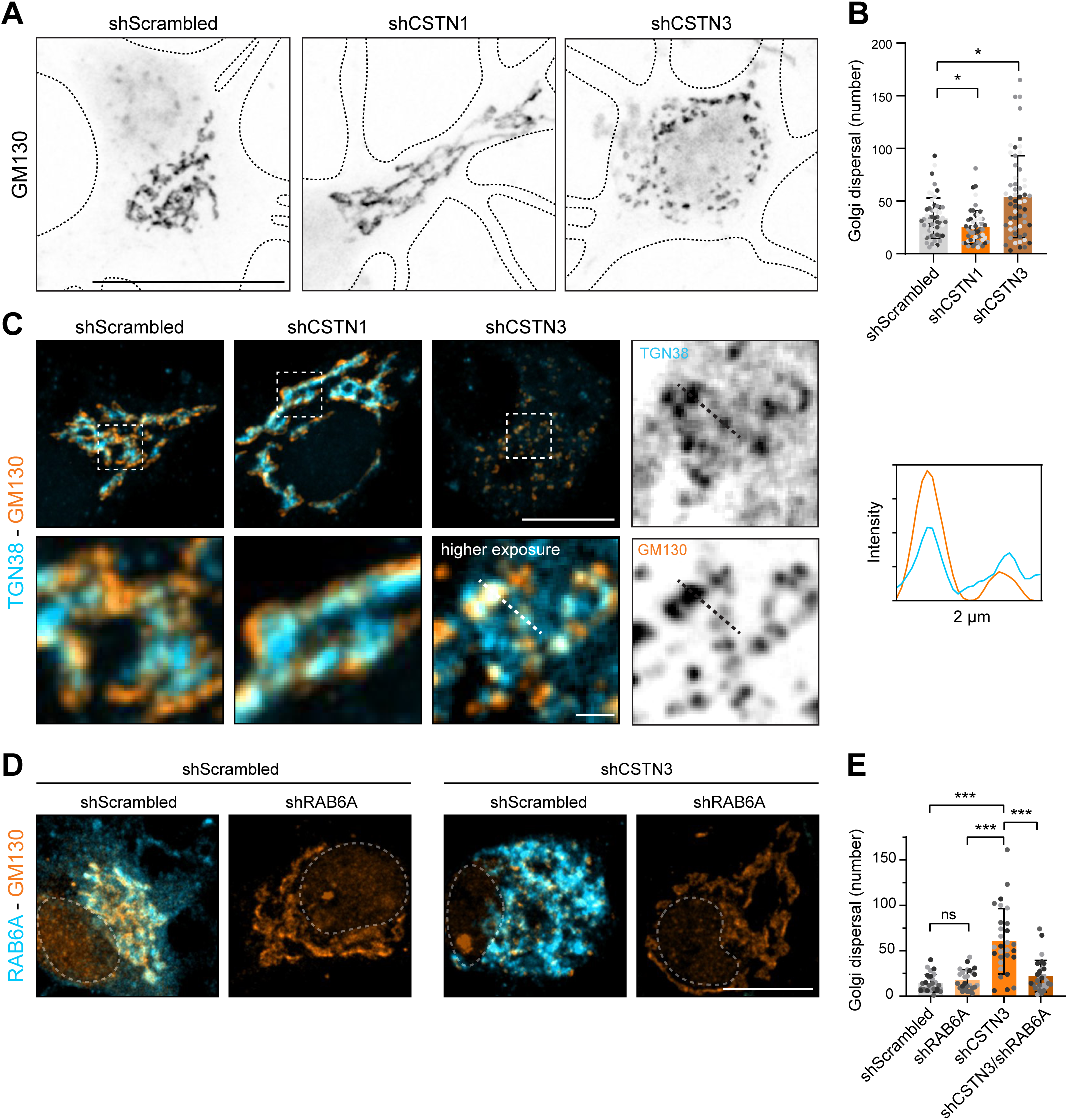
CSTN1 and CSTN3 indirectly regulate Golgi organization through RAB6A. **(A)** Representative images of the soma of neurons transfected at DIV3 with shScrambled-GFP, shCSTN1-GFP or shCSTN3-GFP, stained for GM130 at DIV7. Scale bar: 20 µm. (B) Quantification of the Golgi dispersal as number of Golgi elements. (C) Representative images of the soma of DIV7 neurons transfected at DIV3 with shScrambled-GFP, shCSTN1-GFP or shCSTN3-GFP, stained for GM130 (orange) and TGN38 (blue). Scale bars: 10 µm (overview), 1 µm (zoom). Profile plot shows intensity in arbitrary units. (D) Representative images of the soma of DIV7 neurons transfected at DIV3 with shScrambled-GFP or shCSTN3-GFP, and shRAB6A, stained for GM130 (orange) and RAB6A (blue). Scale bar: 10 µm. (E) Quantification of the Golgi dispersal as number of Golgi elements. All data is presented as mean values ± SD in (C, D). Dots in all dot plots indicate individual cells and replicates are indicated by different colors. In (B, E), Kruskal Wallis followed by Dunn’s correction was used. *p value < 0.05, **p value < 0.01, ***p value < 0.001, ns = not significant.

To further investigate Golgi architecture, we performed double-labeling of GM130 and the TGN38 (Fig. 8C). In both control and CSTN1-depleted neurons, the GM130 and TGN38 signals were closely aligned (Fig. 8C), reflecting the typical cis–trans arrangement of a Golgi stack (van Bommel et al., 2023). Like GM130, TGN38 presented as elongated tubules in CSTN1-depleted neurons. In contrast, in CSTN3-depleted neurons, TGN38 appeared as puncta, which were markedly dispersed throughout the soma but remained co-distributed with GM130 (Fig. 8C), indicating preserved cis–trans organization. This observation is consistent with our EM analyses, where we observed intact Golgi stacks following depletion of either CSTN1 or CSTN3 (Fig. 7E). Together, these findings suggest that loss of CSTN1 or CSTN3 in neurons impairs Golgi ribbon structure, but preserves Golgi stack morphology.

Since we observed increased RAB6A signal in the soma in CSTN3-depleted cells, we wondered whether the effect of CSTN3 in Golgi dispersal could be reversed by RAB6A depletion. Knockdown of RAB6A in control cells, including its splicing variant RAB6A’, did not cause any change in the GM130 staining pattern (Fig. 8D, E). Of note, while RAB6A and RAB6A’ are efficiently knocked down (Fig. S6C), neurons additionally express brain-enriched isoform termed RAB6B (Zhang et al., 2024). However, simultaneous depletion of RAB6A/A’ and CSTN3 reversed the Golgi dispersal as seen upon CSTN3 depletion alone (Fig. 8D, E), indicating that RAB6A/A’ mediate downstream Golgi dispersal in CSTN3-depleted neurons.

To summarize, CSTN1-depleted neurons displayed compact and elongated Golgi cisternae. In contrast, CSTN3-depleted neurons showed dispersal of Golgi stacks throughout the soma, while preserving the cis-trans interface of the Golgi network. Interestingly, RAB6A/A’ depletion rescued the dispersal of Golgi stacks in CSTN3-depleted neurons, indicating that CSTN3 is required for RAB6A/A’ functioning at the Golgi complex.

## Discussion

CSTNs play a critical role in neurodevelopmental processes such as axonal outgrowth and synaptogenesis (Ponomareva et al., 2014; Um et al., 2014), yet their mechanisms of action remain unclear. Here, we dissected the role of the three CSTN paralogs in biosynthetic protein trafficking. We found that depletion of CSTN1 or CSTN3, but not CSTN2, disrupted TGN exit of axonal biosynthetic proteins, impairing RAB6A-positive vesicle delivery to the axon. Interestingly, RAB6A levels at the TGN were differentially affected by CSTN1 and CSTN3 depletion (reduced and increased, respectively), which had opposite effects on Golgi morphology. Our data are summarized in (**Fig. 9**). Based on these data, we propose a model in which CSTN1 and CSTN3 coordinately control cargo delivery to the axon by regulating RAB6A-mediated vesicle budding at the TGN. Our studies provide a foundation to understand the role of CSTNs in axonal development and maintenance.

**Fig. 9.**
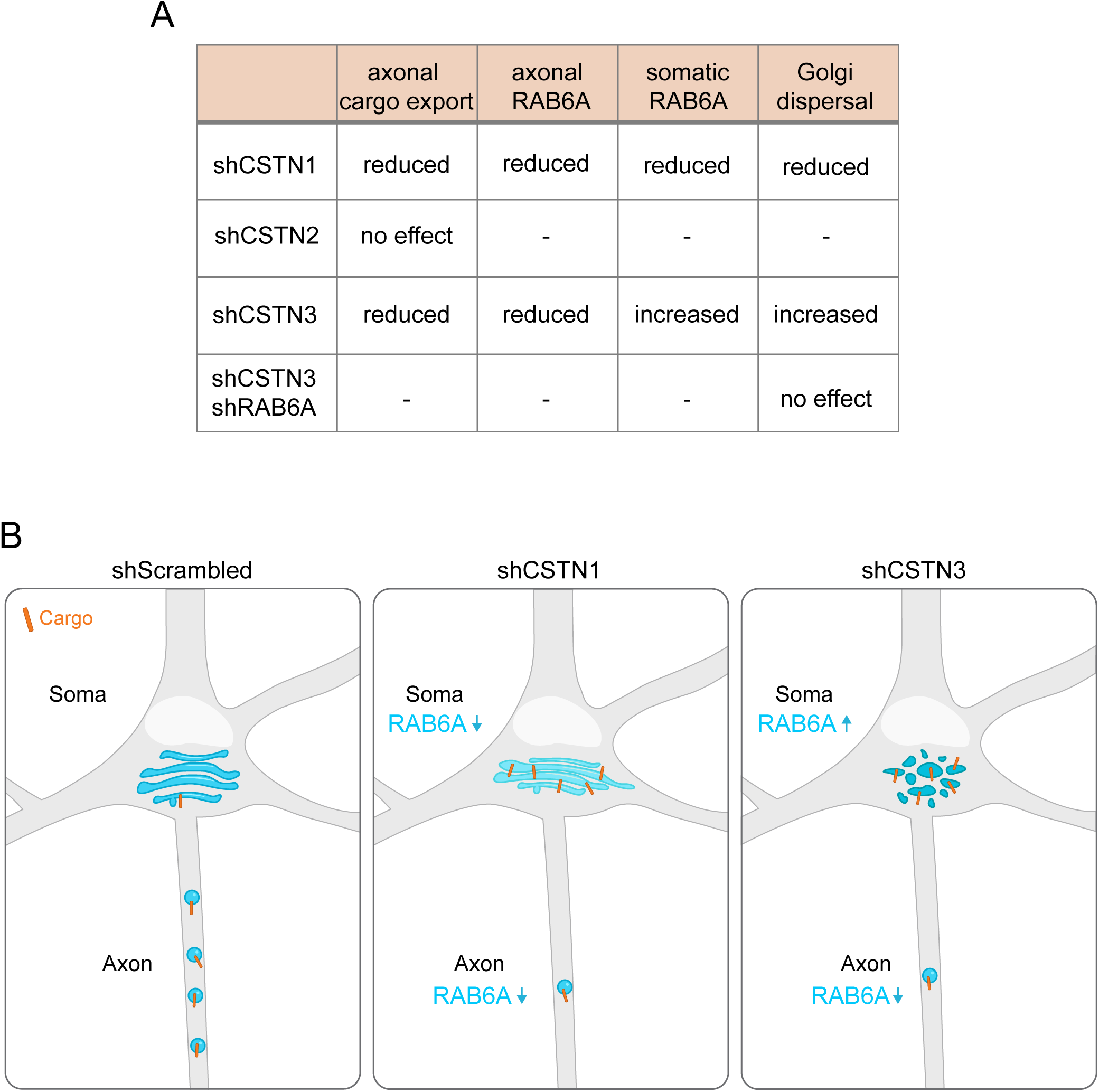
Summary of the effects of CSTN knockdown in neurons. (A) Table summarizing the effects of CSTN1-3 knockdown and combined CSTN3 and RAB6A knockdown on axonal cargo export, axonal RAB6A, somatic RAB6A and Golgi dispersal. (B) Schematic summarizing the effects of CSTN1 and CSTN3 knockdown on the phenotypes in A.

While CSTN1 localization has been inconclusively reported to occur in both in axons and dendrites (Vogt et al., 2001; Hintsch et al., 2002; Konecna et al., 2006), studies on CSTN2 and CSTN3 distribution are limited. To study their endogenous localization in developing primary hippocampal neurons, we applied CRISPR-mediated tagging (Zhong et al., 2021; Droogers et al., 2022). We found that all CSTNs were enriched in the TGN as well as axonal vesicles, consistent with prior evidence for CSTN1 (Konecna et al., 2006; Ludwig et al., 2009). CSTN2 levels appeared lower than that of its paralogs, possibly reflecting its restricted expression to specific neuronal subpopulations (Hintsch et al., 2002; de Ramon Francàs et al., 2017). Despite their predominant axonal enrichment, CSTNs were not entirely excluded from dendrites, which could align with their proposed function as postsynaptic adhesion molecules (Um et al., 2014; Liu et al., 2022). Overexpression of fluorescently tagged CSTN constructs revealed that all three paralogs predominantly traveled in the anterograde direction along the axon, consistent with their proposed interactions with kinesin-1 (Konecna et al., 2006; Araki et al., 2007). CSTN3 showed fewer anterograde transport tracks compared to CSTN1 and CSTN2, which is consistent with its lower binding affinity to kinesin-1, as compared to the other 2 paralogs (Konecna et al., 2006). This may be explained by the fact that CSTN1 and CSTN2 each contain two KBS, while CSTN3 only possesses a single KBS (Konecna et al., 2006).

Overexpressed CSTN constructs largely co-localized on the same transport carriers, suggesting a shared itinerary along the biosynthetic pathway. Although CSTNs are present on the same axonal transport carriers, they may not interact. CSTN3 can form tetramers, but it does not form heterophilic interactions with the other paralogs (Lu et al., 2014). Nevertheless, we found that trafficking of the distinct CSTNs is interdependent. Depletion of CSTN3 reduced the number of CSTN1-positive compartments along the axon, while depletion of CSTN1 led to a subtle increase in CSTN3-positive axonal compartments. The latter was accompanied by a general increase in CSTN3 levels, suggesting CSTN3 expression as a compensatory mechanism for the loss of CSTN1. Intriguingly, previous work indicates that in the absence of CSTN3, the levels of CSTN1 and CSTN2 do not change (Um et al., 2014), suggesting that CSTN3 could – at least partly - compensate for loss of CSTN1 or CSTN2, but not vice versa. This may explain the stronger trafficking defects seen upon CSTN3 depletion.

While CSTN1 is known to regulate the transport of APP (Ludwig et al., 2009; Steuble et al., 2012; Vagnoni et al., 2012), it remained unknown whether the other two CSTN paralogs also contribute to APP trafficking. For this reason, we studied the roles of all three CSTNs in APP transport. Depletion of CSTN1 or CSTN3 reduced axonal APP levels, whereas loss of CSTN2 had no effect, which is consistent with its low expression levels. Reduction of axonal APP upon CSTN1 or CSTN3 depletion was accompanied by APP accumulation in the Golgi region, indicating impaired TGN exit of biosynthetic APP. Beyond APP, we also revealed that all CSTNs colocalized with a diverse set of biosynthetic axonal cargoes that are involved in signaling, cell adhesion and lysosomal functioning, indicating a general role in the trafficking of biosynthetic axonal proteins. Selection of cargo into axonal vesicles could be mediated by interaction of CSTNs with adaptor proteins recognizing signals in cytoplasmic tails of membrane proteins, such as AP-3 (Li et al., 2016). Alternatively, the two cadherin domains in the luminal side of CSTNs could interact with cadherin domains in cargo proteins, such as adhesion molecules (Pettem et al., 2013). Combined, this would equip CSTNs with the unique ability to sort cargo at the TGN by both cytosolic and luminal domains.

To further dissect the role of CSTNs in biosynthetic trafficking, we investigated the effects of CSTN depletion on RAB6A, which marks a broad range of biosynthetic vesicles and plays a key role in cargo packaging at the TGN (Grigoriev et al., 2007; Storrie et al., 2012; Zahavi et al., 2021; Li et al., 2024). Depletion of CSTN1 and CSTN3 reduced the number of RAB6A-positive vesicles in the axon, which aligns with the reduction in transport of various biosynthetic cargoes in these conditions. In addition, ultrastructural analyses by EM revealed a decrease in Golgi-derived vesicles in the soma of CSTN1 and CSTN3 depleted neurons. Although colocalization between CSTN1 / CSTN3 and RAB6A was generally high (70%), we also observed axonal CSTN-positive vesicles that did not contain RAB6A. This population could correspond to retrograde trafficking endocytic compartments transporting CSTNs back to the soma, which are potentially destined for recycling or degradation. Notably, while CSTN1 depletion increased the number of CSTN3 compartments in the axon, it decreased RAB6A abundance, indicating that CSTN3-positive carriers can form and travel also independently of RAB6A.

Intriguingly, we found that depletion of CSTN1 and CSTN3 exerted distinct effects on RAB6A levels at the TGN. Specifically, CSTN1 depletion reduced RAB6A localization at the TGN, whereas CSTN3 depletion increased RAB6A levels at the TGN. This led us to ask whether CSTN1 and CSTN3 could recruit RAB6A via direct interactions. However, RAB6A-GST pulldown experiments did not show direct binding of RAB6A to either CSTN1 or CSTN3, indicating that CSTN1 and CSTN3 indirectly affect RAB6A localization at the TGN. Previous studies have shown that the recruitment of RAB6A to the TGN depends on its GTP active state (Matsuto et al., 2015). Possibly, the reduced total expression levels of RAB6A in CSTN1-depleted cells may be caused by excessive inactive RAB6A-GDP, which is targeted for proteasomal-mediated degradation (Takahashi et al., 2019). In contrast, the increased RAB6A levels at the TGN upon CSTN3 depletion could be due to excessive active RAB6A-GTP, which is recruited to the TGN. In this scenario, CSTNs may act upstream of RAB6A to regulate its activity state. Previous work identified the RIC1 / RGP1 complex as a GEF for RAB6A (Siniossoglou et al., 2000; Pusapati et al., 2012). However, the mechanism by which this complex is recruited to the TGN, and whether CSTNs contribute to this process, remains unclear. It is possible that RAB6A GTP / GDP switching is required for RAB6A-mediated vesicle biogenesis at the TGN, as depletion of either CSTN1 or CSTN3 both reduced vesicle budding from the TGN. More research is required to address whether and how CSTNs regulate the RAB6A GTP / GDP cycle, its recruitment to the TGN, as well as to elucidate how depletion of CSTN1 or CSTN3 leads to these opposite effects.

CSTN1 and CSTN3 depletion also revealed opposite effects on Golgi organization. CSTN1 depletion displayed longer, compact Golgi stacks, while CSTN3 depletion induced Golgi dispersal. Interestingly, RAB6A depletion in non-neuronal cell lines also results in elongated cisterna (Storrie et al., 2012). As depletion of CSTN1 leads to reduced RAB6A levels, it is likely that the cisternal elongation observed in these cells is a consequence of diminished RAB6A levels. Conversely, CSTN3 depletion induced dispersal of the Golgi markers GM130 and TGN38, while retaining an intact cis-trans interface. Overexpression of a RAB6-GTP-locked mutant has been shown to cause a similar phenotype of Golgi dispersal (Martinez et al., 1997), suggesting that the Golgi dispersal in CSTN3-depleted neurons might be caused by increased levels of RAB6. Consistent herewith, we found that RAB6A depletion in CSTN3-depleted cells rescued Golgi dispersal. In addition, depletion of GM130 is also known to disrupt the Golgi ribbon (Puthenveedu et al., 2006), which is consistent with the reduction in GM130 intensity in CSTN3-depleted neurons. Together, these findings indicate distinct, yet intersecting roles for CSTN1 and CSNT3 in Golgi positioning, possibly via RAB6A and / or GM130.

Multiple roles have been proposed for CSTNs, including synaptogenesis and axonal branching (Ponomareva et al., 2014; Um et al., 2014). Potentially, some of these roles can be attributed to the general role of CSTNs in cargo trafficking at the TGN and into the axon, as described in this current paper. However, despite the pronounced effects on axonal transport in CSTN-depleted cells, CSTN triple-knockout mice remain viable (Mori et al., 2022), suggesting the existence of compensatory mechanisms that mitigate loss of CSTN function in vivo.

In summary, CSTN1 and CSTN3 play essential, distinct roles in directing protein exit from the TGN and subsequent axonal trafficking. Their opposing effects on RAB6A levels at the TGN suggest a coordinated regulatory mechanism, where a precise balance of CSTN1 and CSTN3 is required for axonal vesicle biogenesis at the TGN. These findings reveal an unprecedented role for CSTNs in the regulation of polarized protein secretion, which is an essential process for neuronal development and maintenance.

## Declaration of generative AI use

Generative AI (ChatGPT) was used to edit language and improve readability of the manuscript. The authors reviewed and revised all content for scientific accuracy and take full responsibility for the final version.

## Acknowledgement

We thank Anna Akhmanova (Utrecht University, NL) for her valuable suggestions and help with experimental design. We thank the Klumperman lab and Farías lab members and in particular Ha Nguyen and Mai Dan Nguyen for fruitful discussions and assistance with the qPCR experiments. We thank Elena Radul (Utrecht University, NL) for the assistance with protein purification. We thank Wouter Droogers (Radboud University, NL) and Harold MacGillavry (Vrije Universiteit Amsterdam, NL) for the assistance with knock-in generation. We thank René Scriwanek for the assistance with the preparation of the EM figures.

## Data availability

Raw data of all quantifications are available in the Data File. Any other data reported in this paper will be shared by the lead contact upon reasonable request.

## Figure legends

**Fig. S1.**
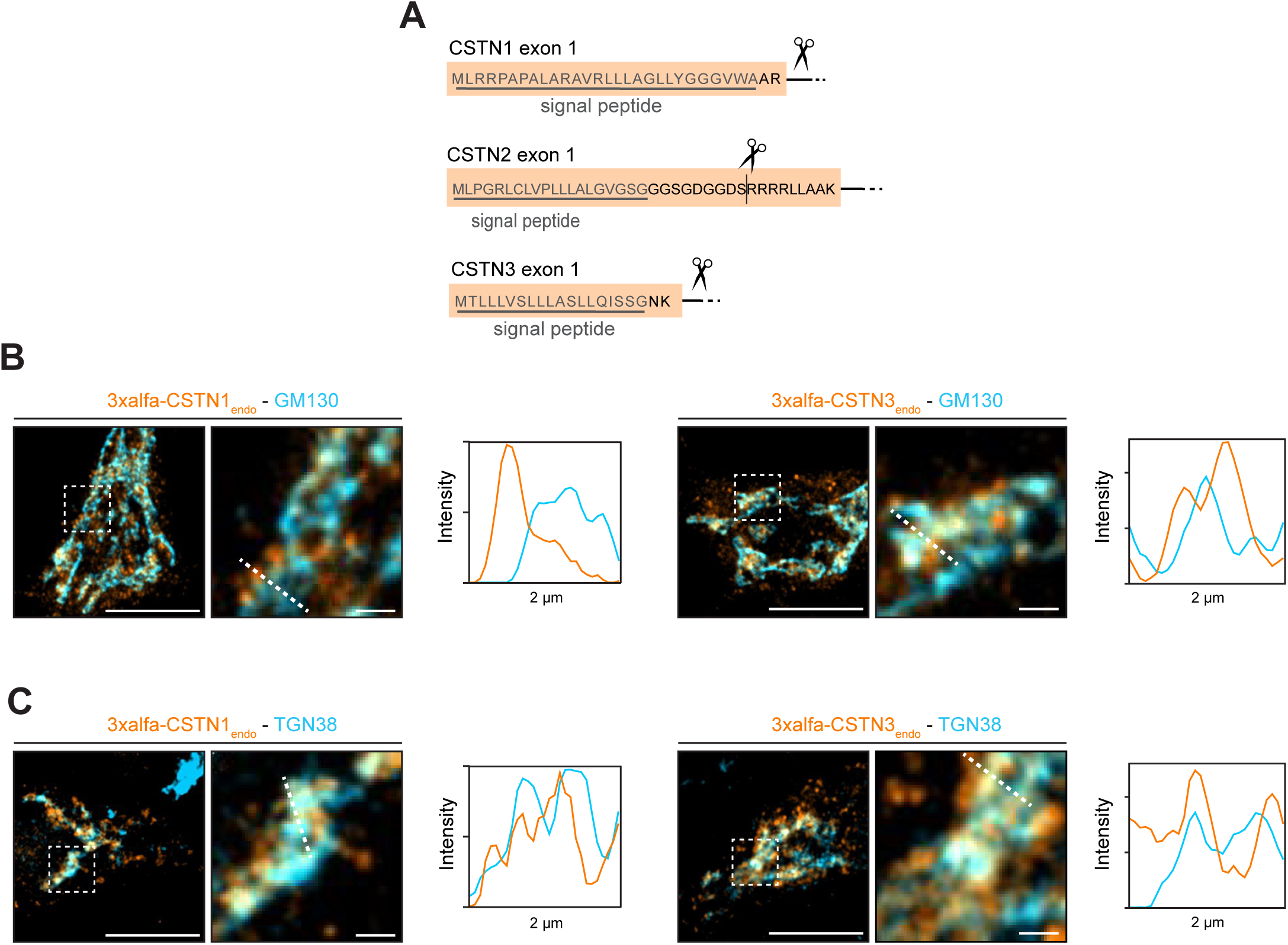
Localization of CSTN1 and CSTN3 to the Golgi. (A) schematic of the Cas9 cutting sites for each CSTN knock-in. (B-C) Confocal images of DIV9 hippocampal neurons with endogenously tagged CSTN1 or CSTN3 (orange), immunolabeled for GM130 (B) or TGN38 (C) (blue). Boxed regions indicate zoom region. Profile plots of indicated dotted lines show intensity in arbitrary units. Scale bars: 10 µm (overview), 1 µm (zoom).

**Fig. S2.**
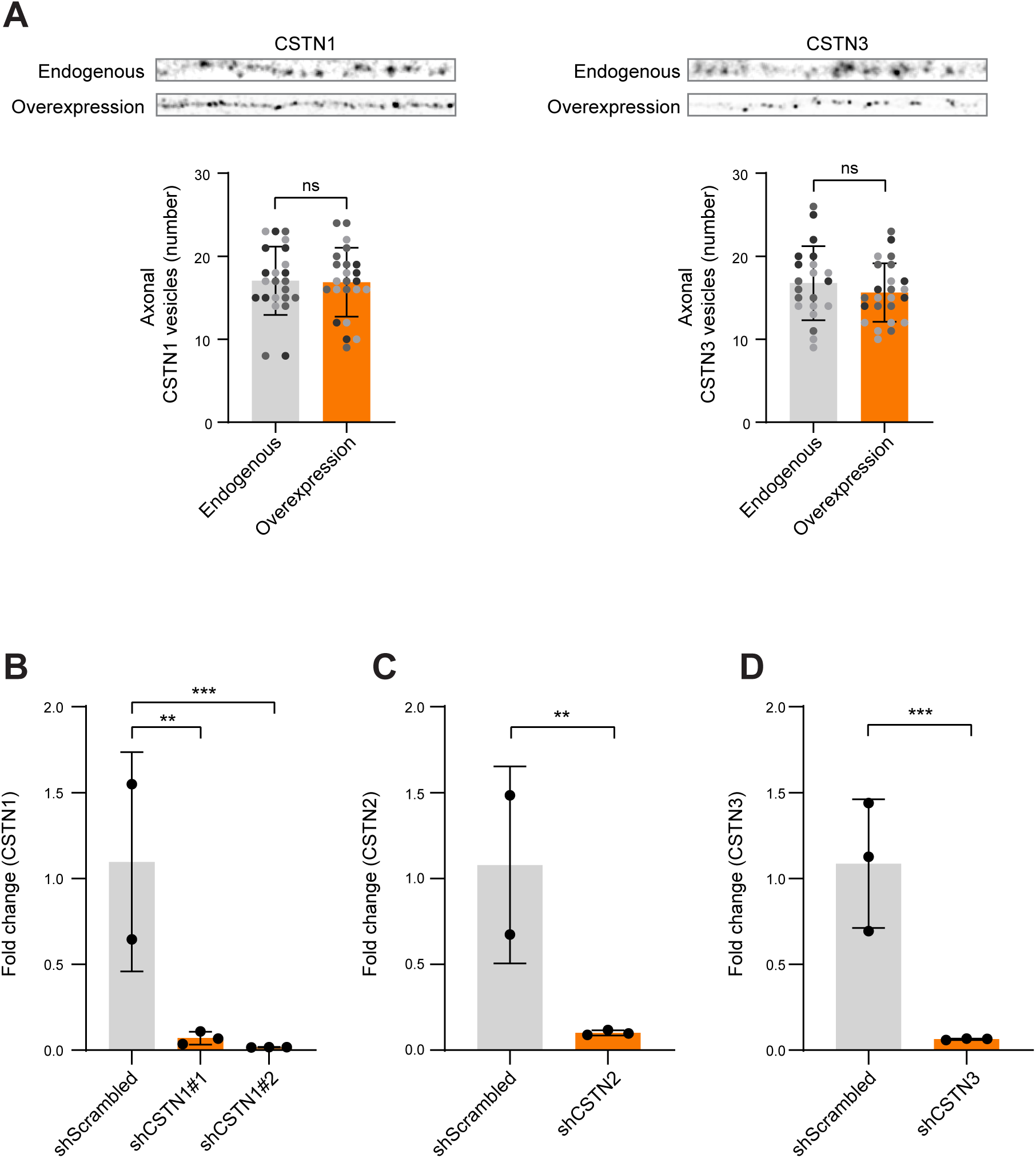
Overexpression of CSTNs do not affect the number of CSTN-positive vesicles along the axon. (A) Representative images of axons (30 µm) from DIV8-9 hippocampal neurons transfected on DIV4 either with endogenous CRISPIE module (top) or HaloTag-CSTN overexpression construct (bottom). Graphs below indicate number of puncta per 30 µm axon. Data for endogenous CSTN1 and CSTN3 were re-used in Fig. 2E and Fig. 2H, respectively. (B-D) Fold change of qPCR of CSTN1 (B), CSTN2 (C) or CSTN3 (D). Cortical neurons were transduced with shRNA on DIV4 and RNA was harvested on DIV8. Fold change data are calculated compared to mean of control. All data is presented as mean values ± SD. Dots indicate individual cells and replicates are indicated by different colors in (A). Dots represent technical replicates (B-D). In (A both graphs, C, D), unpaired t test was used. In (B), ordinary one-way ANOVA followed by Dunnett’s correction was used. Significance was calculated using Ct values in (B-D). *p value < 0.05, **p value < 0.01, ***p value < 0.001, ns = not significant.

**Fig. S3.**
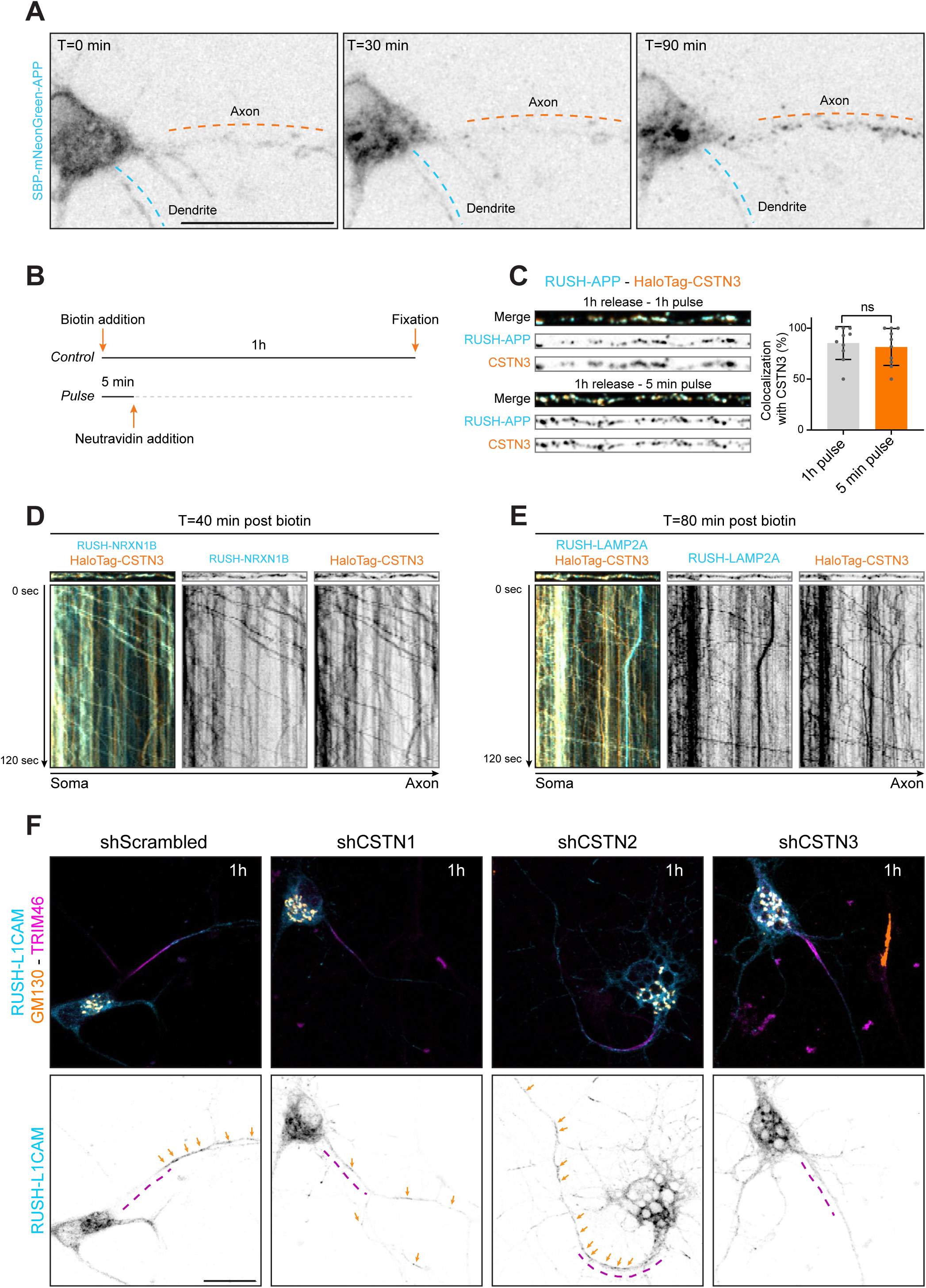
CSTNs colocalize with multiple biosynthetic axonal cargoes. (A) Representative stills of DIV8 hippocampal neurons transfected with RUSH-APP on DIV7. Scale bar: 20 µm. (B) Overview of experimental timeline. Control neurons were exposed for 1h to 100 µM biotin. Neurons in pulse group were exposed for 5 min to 100 µM biotin, followed by addition of 0.3 mg/mL NeutrAvidin. All neurons were fixated 1h post biotin addition. (C) Representative crops of DIV8 hippocampal neurons transfected on DIV7 with HaloTag-CSTN3 and RUSH-APP. Neurons were either continuous exposed to biotin (top) or received a short pulse (bottom). Graph displaying colocalization of RUSH-APP puncta containing CSTN3 (right). (D, E) Representative still images and kymographs of live axons (30 µm) of DIV8 hippocampal neurons transfected on DIV7 with HaloTag-CSTN3 and RUSH-NRXN1B or RUSH-LAMP2A. Kymographs were obtained from 2-minute videos (1s/frame). (F) Representative images of DIV7 hippocampal neurons transfected on DIV3 with RUSH-L1CAM (blue) with shScrambled-GFP, shCSTN1-GFP, shCSTN2-GFP or shCSTN3-GFP and immunolabelled for GM130 (orange) and TRIM46 (magenta) (top). Single channel of RUSH-L1CAM below. Arrows (orange) indicate axonal biosynthetic vesicles. Dashed line (magenta) indicates the AIS. Times indicated in figures indicate time post biotin addition. All data is presented as mean values ± SD. Dots indicate individual cells. In (C), unpaired t test was used. *p value < 0.05, **p value < 0.01, ***p value < 0.001, ns = not significant.

**Fig. S4.**
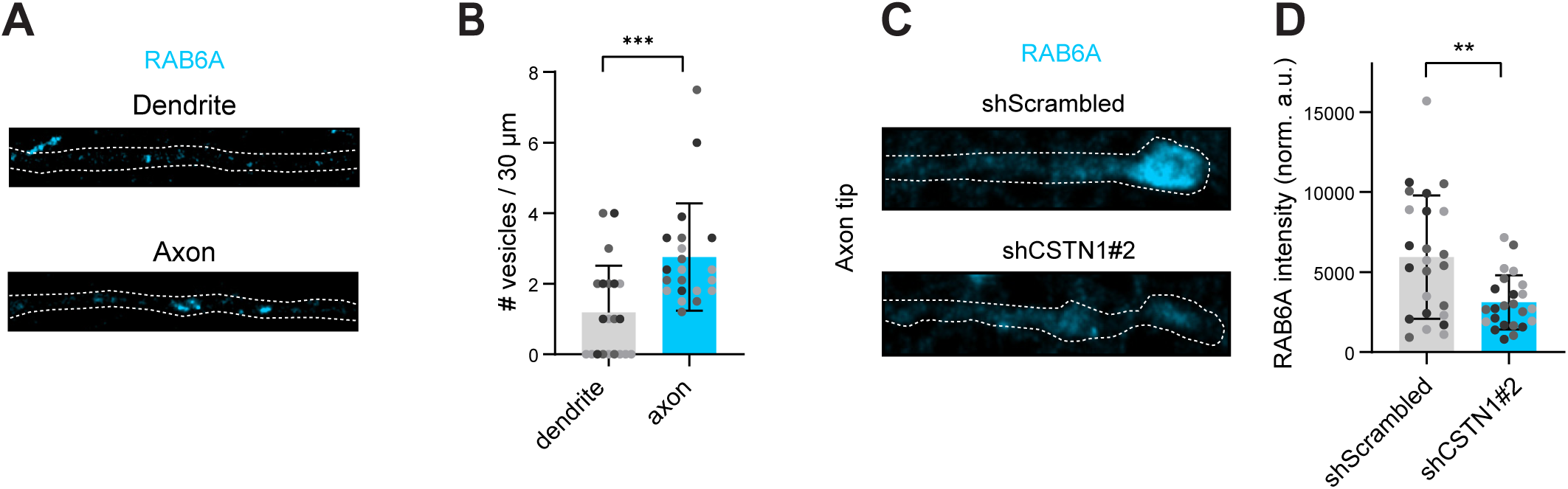
RAB6A-positive vesicles are more abundant in the axon than in dendrites. (A) Representative images of dendrites and axons (30 µm) of DIV7 neurons transfected at DIV3 with shScrambled-GFP and stained for RAB6A. (B) Number of RAB6A-positive puncta in 30 µm of dendrite or axon. For the axon, we normalized the count from Fig. 6D to 30 µm. (C) Representative images of axon tips (20 µm) of DIV8 neurons transfected at DIV4 with shScrambled-GFP or shCSTN1#2-GFP and stained for RAB6A. (D) RAB6A intensity (normalized arbitrary units) measured in the most distal region of the axon tip. In (B), Mann-Whitney test was used. In (D), Welch’s t-test was used. *p value < 0.05, **p value < 0.01, ***p value < 0.001, ns = not significant.

**Fig. S5.**
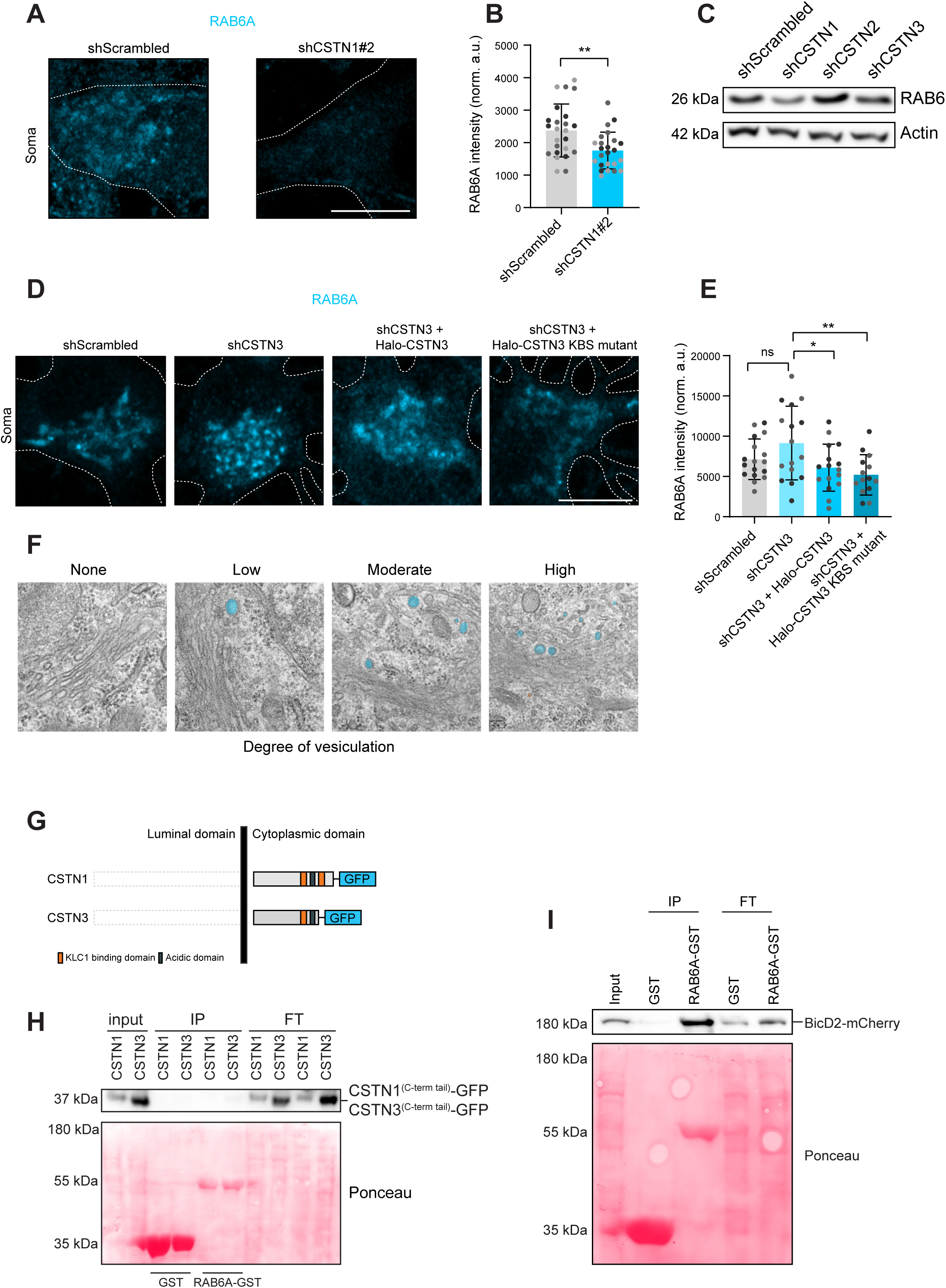
CSTN1 and CSTN3 conversely regulate RAB6A levels in the soma. (A) Representative images of the soma of DIV8 neurons transfected at DIV4 with shScrambled-GFP or shCSTN1#2-GFP and stained for RAB6A. Scale bar: 10 µm. (B) RAB6A intensity (normalized arbitrary units) measured in the perinuclear region. (C) Cortical neurons transduced on DIV4 with lentivirus containing shScrambled-GFP, shCSTN1-GFP, shCSTN2-GFP or shCSTN3-GFP, lysed on DIV8 for western blot for RAB6A (26 kDa) and actin (42 kDa). (D) Representative images of soma of DIV8 neurons transfected on DIV4 with shScrambled-GFP or shCSTN3-GFP with Halo-CSTN3 or Halo-CSTN3 KBS mutant, stained for RAB6A. Scale bar: 10 µm. (E) RAB6A intensity (normalized arbitrary units) measured in the perinuclear region. (F) Visualization of categories for scoring vesiculation at the TGN side of the Golgi complex for Fig. 7F. Vesiculation was scored according to the following scale: none (0%), low (0–10%), moderate (10–30%), and high (30–100%). Percentages represent the proportion of the area occupied by Golgi-derived vesicles, including both clathrin-coated and non-coated structures. (G) Schematic representation of C-terminal CSTN1-3-GFP constructs. (H-I) GST and RAB6A-GST proteins immobilized on beads were incubated with lysates of CSTN1-3-GFP (H) or BicD2-mCherry (I) expressing HEK293T cells. WB was performed using GFP (I) or RFP (H) antibody. 1% input and flowthrough was included in both WB. All data is presented as mean values ± SD. Dots indicate individual cells. In (B), unpaired t-test was used. In (E), ordinary one-way ANOVA followed by Tukey’s correction was used. *p value < 0.05, **p value < 0.01, ***p value < 0.001, ns = not significant.

**Fig. S6.**
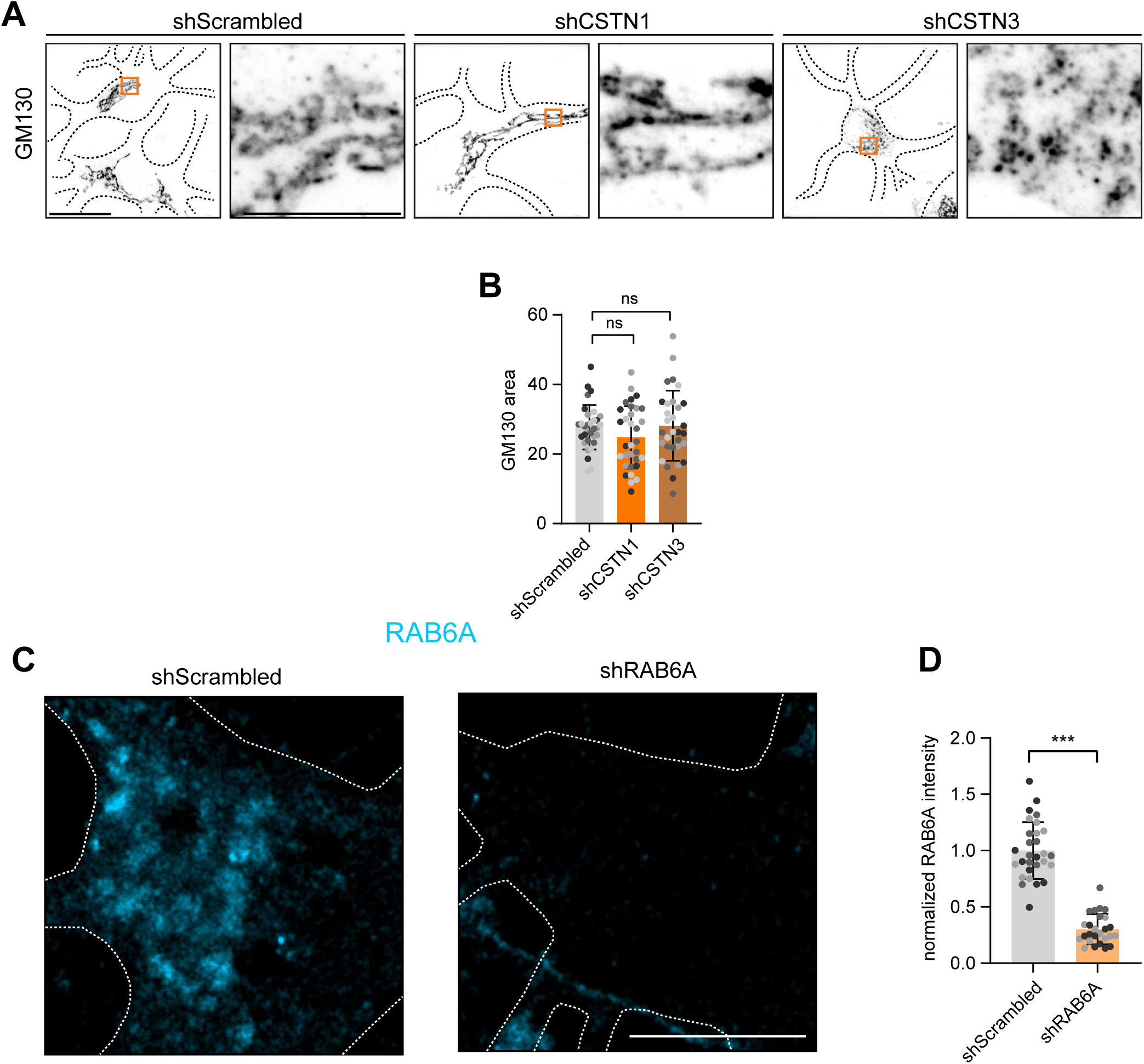
CSTN1 and CSTN3 regulate Golgi organization. (A) Z-projection of expansion microscopy samples from DIV8 hippocampal neurons transduced on DIV4 with shScrambled-GFP, shCSTN1-GFP or shCSTN3-GFP and immunolabelled for GM130. (B) Quantification of total Golgi area of samples shown in Fig. 8A. (C) Representative photos of soma from DIV7 hippocampal neurons transfected on DIV3 with shScrambled or shRAB6A and immunolabelled for RAB6A. Dotted lines indicate cell outline. (D) RAB6A intensity of shScrambled and shRAB6A expressing neurons. Values are expressed as fold change to control. Mann-Whitney test was used. All data is presented as mean values ± SD. Dots indicate individual cells. In (E), ordinary one-way ANOVA test was used followed by Dunnet’s correction. *p value < 0.05, **p value < 0.01, ***p value < 0.001, ns = not significant.

**Video S1**. **CSTN paralogs are transported anterogradely along axonal tracks.** Representative videos (Spinning Disk microscope) from axon crops of DIV7 hippocampal neurons expressing HaloTag-CSTN1, HaloTag-CSTN2 or HaloTag-CSTN3. Neurons were imaged 3 minutes every second.

**Video S2. APP colocalizes with anterograde moving CSTN carriers.** Representative videos (Spinning Disk microscope) from axon crops of DIV7 hippocampal neurons expressing HaloTag-CSTN1, HaloTag-CSTN2 or HaloTag-CSTN3 (orange) together with mNeonGreen-APP (blue). Neurons were imaged 3 minutes every second.

**Video S3. RUSH-APP exits the TGN via CSTN-positive carriers.** Representative videos (Spinning Disk microscope) from HeLa WT cells expressing HaloTag-CSTN1, HaloTag-CSTN2 or HaloTag-CSTN3 (orange) together with RUSH-APP (blue). Neurons were imaged 2 minutes every 1.5 second. Time post biotin addition is indicated in Fig. 4B. Biotin addition took place at T=0.

**Video S4. Release dynamics of RUSH-APP in hippocampal neurons.** Representative videos (Spinning Disk microscope) from DIV8 hippocampal neurons expressing RUSH-APP following release. Neurons were imaged 90 minutes every minute. Biotin addition took place at T=0.

## Materials and methods

### Animals

All experiments were approved by the DEC Dutch Animal Experiments Committee (Dier Experimenten Commissie), performed in line with the guidelines of University Utrecht and in agreement with Dutch law (Wet op de Dierproven, 1996). The animal protocol has been evaluated and approved by the national CCD authority (license AVD10800202216383). Female pregnant Wistar rats were obtained from Janvier, and embryos (both genders) at embryonic (E)18 stage of development were used for primary cultures of hippocampal and cortical neurons. The animals, pregnant females and embryos have not been involved in previous procedures.

### Primary neuron culture and transfection

Hippocampi or cortices from embryonic day 18 rat brains were dissected and dissociated with trypsin for 15 min and plated on coverslips coated with poly-L-Lysine (37.5 μg/mL) and laminin (1.25 μg/mL) at a density of 100,000/well (12-well plates) for (live) imaging or 1,000,000/well (6-well plates) for biochemical assays (*Western Blot*). The day of neuron plating corresponds to day-in-vitro 0 (DIV0). Neurobasal medium (NB) supplemented with 2% B27 (GIBCO), 0.5 mM glutamine (GIBCO), 15.6 μM glutamate (Sigma), and 1% penicillin/streptomycin (GIBCO) was used to maintain neurons incubated under controlled temperature and CO2 (37 °C, 5% CO_2_).

Hippocampal neurons were transfected using Lipofectamine 2000 (Invitrogen). Briefly, DNA was mixed with 1.2 - 3.3 μL of Lipofectamine 2000 in 200 μL NB medium containing no supplements or 0.5 mM glutamine and incubated for 20 min at room temperature. DNA mix was subsequently added to neurons and incubated for 1 h (37 °C, 5% CO2). Next, neurons were washed with NB and transferred to half of their original medium and half fresh supplemented NB at 37 °C in 5% CO_2_ until fixation or imaging at different days in vitro (DIV) as indicated in figure legends.

For RUSH assays, biotin was removed from B27 (GIBCO) using Zebra^TM^ Dye and Biotin Removal Spin Columns (Invitrogen) according to standard manufacturer’s protocol. Following transfection, neurons were stored in supplemented NB medium containing biotin free B27 at 37 °C in 5% CO_2_ until fixation or imaging at different days in vitro (DIV) as indicated in figure legends.

### Culture of cells lines and transfection

HeLa parental cells were obtained from DSMZ (ACC 57). HEK293T cells were obtained from ATCC and were a kind gift from Harold MacGillavry. Cell lines were maintained in DMEM (HPSTA; Capricorn) supplemented with 10% FBS and 1% penicillin and streptomycin. Cells were kept at 37 °C and 5% CO2. HeLa and HEK293T cells were passaged twice or thrice per week respectively after reaching 70-90% confluence. Cells were transfected after reaching 70% confluence. For live-cell microscopy, cells were transfected with Lipofectamine 2000 (ThermoFisher Scientific). In brief, DNA was mixed with 2 μL of Lipofectamine 2000 in 25 μL optiMEM (GIBCO) containing no supplements and incubated for 20 min at room temperature. DNA mix was subsequently added to cells and incubated for 2-3 h (37 °C, 5% CO2). Following incubation with DNA mix, medium was replaced to supplemented DMEM and incubated ON (37 °C, 5% CO2) until imaging.

### Lentivirus packaging and transduction

HEK293T cells were transfected with shRNA-encoding vector, packaging vector psPAX2 and envelope vector pMD2.G with a ratio of 3:2:1 using PEI Max (3:1 PEI Max:DNA ratio). Transfection mix was prepared in optiMEM (GIBCO) incubated 20-min at room temperature. DNA mix was subsequently added to HEK293T cells and incubated for approximately 6 h (37 °C, 5% CO_2_). Medium was removed after 6 h and changed to NB supplemented with 0.5 mM glutamine. Supernatant from packaging HEK293T cells was collected 48 to 72 h after transfection and filtered through a 0.45 μm filter. Supernatant was further concentrated using an Ultra Centrifugal Filter 100 kDa MWCO (AMICON) at 4 °C 1,000 g for 15 to 30 min to a final volume of ±100 μL. Day of lentiviral transduction is indicated in figure legends. 1-2 μL/well (12-well plate) or 10-20 μL/well (6-well plate) of concentrated virus was used. Respective days of lentiviral transduction are indicated in figure legends.

### DNA and shRNA constructs

The following plasmids were used in this study:

FUGW was a gift from David Baltimore (Addgene plasmid #14883) (Lois et al., 2002). psPAX2 and pMD2.G were gifts from Didier Trono (Addgene plasmids #12260 and #12259). pGW1-mCherry and pSuper were a gift from Juan Bonifacino. BicD2–mCherry, pGEX_GST and pGEX_RAB6-GST were a gift from Anna Akhmanova (Matanis et al., 2002). FUGW_Strep-KDEL_SBP-HA-NF186-mNeonGreen was a gift from Ha Nguyen (Farias lab). pLKO.1 puro was a gift from Bob Weinberg (Addgene plasmid #8453) (Stewart et al., 2003). Halo-TfR-RUSH was a gift from Jennifer Lippincott-Schwartz (Addgene plasmid #166905) (Weigel et al., 2021). pEGFP-n1-APP was a gift from Zita Balklava & Thomas Wassmer (Addgene plasmid #69924) (Currinn et al., 2015). pOC1 FLAG-SpCas9 was a gift from Harold MacGillavry (Addgene #183423). FUGW_SBP-mNeonGreen-APP and mNeonGreen-APP were generated previously (Draper et al., 2026). FUGW_RUSH, FUGW_StrepKDEL_SBP-LAMP2A-mNeonGreen and FUGW_StrepKDEL_SBP-SYT1-mNeonGreen were previously generated (Li et al., 2024).

Plasmids generated for this study:

To generate the CSTN knock-in constructs, we applied the pORANGEtrap system developed by the MacGillavry group (unpublished). ORANGEtrap destination vector and 3xalfa donor plasmids with appropriate frameshift corrections were gifts from Harold MacGillavry. Target sequences for CSTN1, CSTN2 and CSTN3 were selected using CRISPOR (Concordet and Haeussler, 2018). CSTN1 (intron 1): ACACTCCTAAGGAATCAGC, CSTN2 (exon 1): GCGACGGCGGGGACAGCCGG, CSTN3 (intron 1): GAGCTAGGAGGACCATCCTG. These target sequences were incorporated into the oligos in table 1. Golden Gate cloning with Sap1 restriction enzyme (Thermofisher) was used to incorporate the annealed oligos and the 3xalfa donor sequence into the ORANGEtrap destination vector, in a similar fashion as the SapTrap method (Schwartz and Jorgensen, 2016). Of note, for CSTN1 and CSTN3, the donor sequence is inserted into the first intron by CRISPIE method (Zhong et al., 2021), while the donor sequence is inserted into the first exon of CSTN2 using the ORANGE method, see figure 1B, 1C and Suppl. Fig 1A (Willems et al., 2020; Droogers et al., 2022).

pEF1_HaloTag-CSTN template constructs were ordered from VectorBuilder. HaloTag-CSTN was subsequently amplified by PCR from pEF_HaloTag-CSTN and transferred to pEGFP-N1 digested with XhoI and AgeI using Gibson Assembly. For the generation of pCMV_mNeonGreen-CSTN1 and pCMV_mNeonGreen-CSTN1 constructs, CSTN sequences were amplified from pCMV_HaloTag-CSTN. mNeonGreen was amplified from BACE1-mNeonGreen (Draper et al., 2026). All fragments were inserted using Gibson Assembly. KBS mutation (W897A) was generated in pCMV_HaloTag-CSTN3 by using the QuickChange II XL Site-Directed Mutagenesis Kit. shResistant HaloTag-CSTN constructs (HaloTag-CSTN3 and HaloTag-CSTN3 KBS mutant) were generated by restriction of FUGW (Addgene plasmid #14883) (Lois et al., 2002) with NheI and XbaI and subsequent insertion using Gibson Assembly of shResistent PCR fragments. CSTN1_C-term_–GFP, CSTN3_C-term_–GFP were generated by restriction of FUGW (Addgene plasmid #14883) (Lois et al., 2002) with NheI and XbaI. GFP was amplified from pEGFP-N1-APP (Addgene plasmid #69924) (Currinn et al., 2015). CSTN fragments were amplified from pCMV_HaloTag-CSTN plasmids. PCR insertion took place by using Gibson Assembly.

For generation of FUGW_Strep-Ii_L1CAM-mNeonGreen-SBP, we first the generated FUGW_Strep-Ii vector. FUGW vector (Addgene plasmid #14883) was digested with BamHI and EcoRI. Strep-Ii-Intron-IRES was amplified from pIRES-neo3-Strep-Ii-LAMP2-SBP-FKBP-GFP (a gift from Judith Klumperman). The resulting PCR product was assembled with digested FUGW using Gibson Assembly to generate FUGW-RUSH-Ii for subsequent cloning purposes. L1CAM was amplified from GW1-PAGFP-mL1CAM (gift from Casper Hoogenraad). mNeonGreen was amplified from FUGW_StrepKDEL_SBP-SYT1-mNeonGreen. A 20 aa linker was added by overhang PCR. SBP was amplified by from FUGW_StrepKDEL_SBP-SYT1-mNeonGreen. FUGW_Strep-Ii was digested with EcoRI and BamHI. Fragments were inserted by Gibson Assembly. For the generation of FUGW_Strep-KDEL_IL2-SBP-TRKB-mNeonGreen, IL2-SBP-TRKB was amplified from pIRES3_IL2-SBP-TRKB-EGFP (Zahavi et al., 2021) and mNeonGreen was amplified from BACE1-mNeonGreen (Draper et al., 2026). FUGW_Strep-KDEL (Li et al., 2024) was digested using BamHI and EcoRI and PCR fragments were inserted by Gibson Assembly.

The pFSW-StrepKDEL vector was generated by amplification of Strep-KDEL-intron-IRES from Str-KDEL_SBP-EGFP-Ecadherin (Addgene plasmid #65286) with XbaI and EcoRI sites introduced and inserted into restricted pFSW-67 (a gift from Harold MacGillavry) by GIBSON assembly. pFSW-StrepKDEL_SBP-TRKB-mNeonGreen (referred to as SBP-TRKB-mNeonGreen) and pFSW-Strep-KDEL_SBP-mNG-APP (referred to as SBP-mNG-APP) were generated by amplification of SBP-TRKB-mNeonGreen from FUGW_Strep-KDEL_IL2-SBP-TRKB-mNeonGreen and amplification of SBP-mNeonGreen-APP from FUGW_Strep-KDEL_SBP-mNeonGreen-APP (Draper et al., 2026). These fragments were inserted into pFSW_StrepKDEL vector digested with EcoRI and BamHI by Gibson assembly. FUGW_StrepIi_NRXN1B_SBP_mNeonGreen was generated by amplifying the NRXN1B signal peptide and NRXN1B from FUGW_RUSH-KDEL-NRXN1B-mScarlet (gift from Ha Nguyen, Farías lab). SBP and mNG were amplified from FUGW_StrepKDEL_SBP-SYT1-mNeonGreen. These fragments were inserted into FUGW_StrepIi digested with EcoRI and BamHI by GIBSON assembly.

The following sequences for rat-shRNAs inserted to pLKO.1-puro and / or pSuper were used in this study: shScramble (5’-ACAATAGCTTACTACCAAT-3’), shCSTN1#1 (5’- GCAAAGAGCATCAGTATAA-3’), shCSTN1#2 (5’-CCATCACGCTTGCAGTTTT-3’), shCSTN2 (5’- TGGAAAGCCAGAAGGTGAT-3’), shCSTN3 (5’-CATTGAGAACACAGAGAAG-3’), shRAB6A (5’-CATCATGCTAGTAGGAAATAA-3’). For the validation of shRNA CSTN target sequences, we transduced rat cortical neurons on DIV4 and harvested total RNA on DIV8 using RNeasy Mini kit (Qiagen, 74104). cDNA was reverse-transcribed using SuperScript III First-Strand Synthesis System (Invitrogen, 18080-051) according to manufacturer’s instructions. Quantification of target genes was done by qPCR using Power SYBR Green PCR Master Mix (Applied Biosystems, 4367659) and ViiA 7 Real-Time PCR System (Applied Biosystems). Data was extracted from ViiaA 7 Real-Time PCR system using QuantStudio Software V1.3 (Applied Biosystems). qPCR oligos are described in Table 1.

#### Validation of shRAB6A

The amount of RAB6A in the soma was quantified by drawing a polygon selection in the perinuclear area and measuring the mean intensity. This value was normalized by dividing each value by the average intensity of the scrambled condition within each replicate.

### Reagents and antibodies

For antibodies used in this study, see table 2. Other reagents used were: Lipofectamine 2000 (Invitrogen, #1639722), Zebra^TM^ Dye and Biotin Removal Spin Columns (Invitrogen), PEI (Polysciences, #19850), biotin (Sigma-Aldrich, #B4501), Gibson Assembly Mix (New England Biolabs, #E2611), Mix-n-Stain CF640R antibody labeling kit (Biotium, #92245), Glutathione Sepharose™ 4B GST-tagged protein purification resin (Cytiva, #17075601), biotin (Sigma-Aldrich, #B4501), acrylamide 40% solution (Sigma-Aldrich, #A4058), N,N′-methylenebisacrylamide solution (Sigma-Aldrich, #M1533), APS (Sigma-Aldrich, #215589), TEMED (Bio-Rad, #1610800), acrylic acid (Sigma-Aldrich, #147230), JFX554 and JFX650 were kindly provided by the Lavis Lab (Janelia).

### Cell lysis and Western Blot

Primary rat cortical neurons were cultured in 6-well plates and washed two times in ice cold PBS before scraping. Cells were scraped and lysed in RIPA buffer (150 mM NaCl, 50 mM Tris-HCl pH 7.4, 0.1% SDS, 0.5% sodium-deoxycholate, 1% Triton X-100 and 1× protease inhibitor (Roche)) followed by incubation for 15 min at 4 °C. Lysates were cleared by centrifuging for 15 min at 16,000 x g 4 °C after which the pellet was discarded. Protein concentration was measured using the Pierce™ BCA Protein Assay Kit (Thermo Fisher Scientific) in order to equalize protein loading on gels. 20 μg of protein was used for all experiments, unless indicated differently in figure legend. Lysates were resolved by SDS-PAGE on a 12% Bis-Acrylamide (Bio-Rad) gel and transferred to a PVDF membrane (Bio-Rad). These membranes were blocked for 1 h at room temperature in TBS-T containing 3% BSA. Primary antibodies were diluted in blocking buffer and incubated ON at 4 °C. After three washes with TBS-T, membranes were incubated with HRP or IRDye secondary antibody in blocking buffer for 1 h at room temperature and washed three times with TBS-T prior to imaging. For samples incubated with HRP secondary antibody: membranes were incubated in Clarity Western ECL Substrate (Bio-Rad) and developed using ImageQuant 800 (AMERSHAM). For samples incubated with IRDye secondary antibody: membranes were developed on an Odyssey CLx imaging system (LICOR) with Image Studio version 5.2.

### Protein purification and GST-pulldown

For in vitro binding studies, recombinant GST and RAB6A–GST fusion proteins cloned in pGEX plasmids were kindly provided by Anna Akhmanova (Matanis et al., 2002). GST and RAB6A–GST were expressed in BL21 E. coli. Protein expression was induced by the addition of 0.15 mM IPTG, followed by overnight incubation at 18 °C. Bacterial cultures were harvested by centrifugation at 4,500 g and the resulting pellets were stored at −80 °C until further use. After thawing, bacterial pellets were resuspended in lysis buffer (PBS containing 20 mM Tris-HCl, 0.1% Tween-20, 250 mM NaCl, 5 mM MgCl₂, 1 mM DTT, 1× cOmplete protease inhibitor). Cells were lysed by sonication using a Soniprep 150 (five cycles of 30 sec on / 30 sec off). Lysozyme was then added to a final concentration of 2 mg/mL and lysates were incubated on ice for 60 min. Lysates were clarified by centrifugation at 27,000 g for 30 min at 4 °C and the supernatant was collected. GST or RAB6A–GST proteins were purified by incubating the clarified supernatant with equilibrated glutathione-Sepharose 4B beads (Cytiva) for 2 h at 4 °C. The resin was subsequently collected by centrifugation at 500 g for 5 min and removal of supernatant. Beads were washed twice with binding buffer (PBS containing 0.1% Tween-20, 250 mM NaCl, 5 mM MgCl₂, 1 mM DTT, 100 µM GDP) and subsequently stored at 4 °C.

For GST pulldown experiments, HEK293T cells (∼8.8 × 10⁶ cells) expressing CSTN1_C-term_–GFP, CSTN3_C-term_–GFP, or BicD2–mCherry (Matanis et al., 2002) were lysed in 1 mL lysis buffer (10 mM Tris-HCl pH 7.5, 150 mM NaCl, 0.5 mM EGTA, 1% NP-40, 5 mM MgCl₂, 0.5 mM CaCl₂, 1× cOmplete protease inhibitor, 100 µM GDP). Prior to incubation with immobilized GST fusion proteins, cell lysates were precleared by incubation with 20 µL equilibrated glutathione-Sepharose 4B beads for 30 min at 4 °C to remove non-specific binding proteins. Lysates were then centrifuged at 500 g for 5 min and supernatant was collected. The precleared lysates were incubated for 1 h at 4 °C with immobilized GST (5 µL beads) or RAB6A–GST (20 µL beads). Following incubation, beads were washed three times with lysis buffer and subsequently subjected to SDS-PAGE as described in section (*Cell Lysis and Western Blot*). For all experiments, 1% input and flow-through fractions were collected and analyzed by Western blot.

### Live-cell imaging

For live-cell imaging of HeLa cells and primary rat hippocampal neurons, cells were imaged on a Nikon Eclipse Ti or Nikon Eclipse Ti2 equipped with an incubator chamber (INUG2-ZILCS0H2; Tokai Hit) on a motorized stage (ASI), a Nikon Plan Apo VC 100× N.A. 1.40 oil objective, and a spinning disk confocal scanner unit (CSU-X1-A1; Yokogawa). Illumination was performed using a CoolLED pE4000 (CoolLED) LED device and ET-EGFP (49002; Chroma), ET-mCherry (49008; Chroma), ET-CY5 (49006; Chroma), and ET-CY5.5 (49022; Chroma) filters. All images were acquired with a Coolsnap HQ2 CCD camera (Photometrics). The microscope was controlled using μManager software (Edelstein et al., 2014). Coverslips were mounted in a metal ring and cells were supplemented in their original medium. Cells were imaged in an incubation chamber that maintains optimal temperature and CO2 (37 °C and 5% CO2). For multicolor imaging, sequential imaging was used with a maximum exposure of 0.3 s per channel. For imaging of constructs containing HaloTag, cells were incubated with either JF554 or JFX650 (100 nM) in NB without supplements prior to imaging (37 °C and 5% CO2). To identify axons, neurons were either co-transfected with a fill and identified by morphology or stained for 15 min with a CF640R-conjugated antibody against the axon initial segment (AIS) protein neurofascin (NF640R). For the RUSH assay using live cells, biotin was added on stage (100 μM) for synchronized release of RUSH cargo. Total time and intervals of imaging acquisition for each experiment are indicated in figure legends.

### Immunofluorescence staining and imaging

Medium was removed from wells, cells were washed with PBS supplemented with calcium and magnesium (PBS-CM) and incubated with prewarmed (37 °C) 4% paraformaldehyde supplemented with 4% sucrose in PBS for 10 min. Cells were permeabilized with 0.2% Triton X-100 in PBS-CM for 15 min, followed by blocking with 0.2% porcine gelatin in PBS-CM for 1 h at room temperature. Alternatively, cells were blocked and permeabilized for 1 h at room temperature with PBS-CM containing 0.05% saponin and 0.2% porcine gelatin for 1 h at room temperature. For samples treated with Triton, primary and secondary antibodies were diluted in 0.2 % porcine gelatin in PBS-CM. For samples treated with saponin, antibodies were diluted in 0.05% saponin and 0.2% porcine gelatin in PBS-CM. Primary antibodies were incubated for 2 h at room temperature or at 4 °C overnight. Secondary antibodies were incubated for 1 h at room temperature. Cells were washed three times in PBS-CM in between all steps. Samples were mounted in Fluoromount-G Mounting Medium (Thermofisher Scientific).

Imaging took place on a confocal laser-scanning microscope (LSM700, with Zen imaging software (Zeiss) version 8.1.7.484) equipped with Plan-Apochromat ×63 NA 1.40 oil DIC and EC Plan-Neofluor ×40 NA1.30 Oil DIC objectives or on a LSM900 (with Zen blue imaging software version 3.7.97.07000 (Zeiss) equipped with Plan-Apochromat ×63 NA 1.40 oil DIC objective).

### Electron microscopy

Neurons were grown in 6 cm dishes and transduced on DIV4 with pLKO-GFP shRNA constructs as described in previous section (*Lentivirus packaging and transduction in neurons*). Cells were fixed on DIV8 by adding 2% FA, 0.2% GA in 0.1 M PHEM buffer (pH 7.4) to an equal volume of medium for 5 min at room temperature. Afterwards, fixative was refreshed and cells were incubated for 2 h. Cells were stored in 1% formaldehyde at 4 °C until further processing. Post-fixation was performed using 1% wt/vol OsO4, 1.5% wt/vol K3Fe(III)(CN)6 in 0.1 M PHEM buffer for 2h at 4°C. Next, cells were dehydrated with ethanol and embedded in Epon (Electron Microscopy Sciences). Ultrathin sections were stained with uranyl acetate and lead citrate using AC20 (Leica). Images were taken on a JEM1010 (JEOL) equipped with a Veleta 2k×2k CCD camera (EMSIS, Munster, Germany)

### Expansion microscopy

Anchoring was performed by incubating fixed neurons (*Immunofluorescence staining and imaging*) in 1.4% (w/v) formaldehyde and 3% (w/v) acrylamide (Sigma-Aldrich, A4058) in PBS for 3 h at 37°C. After several washes with PBS, neurons were incubated in U-ExM gelation solution [19% (w/v) sodium acrylate, 10% (w/v) acrylamide (Sigma-Aldrich, A4058), 0.1% (w/v) N,N′-methylenebisacrylamide (Sigma-Aldrich, M1533), 0.5% (w/v) APS (Sigma-Aldrich, 215589), 0.5% (w/v) TEMED (Bio-Rad, 1610800), and 1× PBS] for 2–3 min on ice, followed by polymerization for 1 h at 37°C (Gambarotto et al., 2019). The 38% (w/v) sodium acrylate stock solution was prepared as previously described (Damstra et al., 2022). Briefly, acrylic acid (Sigma-Aldrich, 147230) was neutralized with 10 M sodium hydroxide to a final pH of 7.5–8 and stored at −20°C. Gels were denatured in a solution containing 14% SDS, 300 mM Tris (pH 9), and 500 mM NaCl for 1.5 h at 95°C. Following denaturation, gels were washed twice for at least 30 min in Milli-Q water, and then twice in PBS for at least 30 min with shaking. The expansion factor used for image calibration was determined by dividing the gel diameter (after repeated washes in water until no further expansion occurred) by the coverslip size. For post-expansion labeling, gels were cut into smaller pieces and incubated overnight at room temperature with primary antibodies diluted 1:250 in PBS supplemented with 2% (w/v) BSA and 0.1% (v/v) Triton X-100, with shaking. Gels were then washed 4 × 15 min in PBS containing 0.1% (v/v) Triton X-100. Subsequently, gels were incubated with secondary antibodies diluted 1:500 in PBS supplemented with 2% (w/v) BSA and 0.1% (v/v) Triton X-100 for 3 hrs at room temperature with shaking, followed by 4 × 15 min washes in PBS containing 0.1% (v/v) Triton X-100. Finally, gels were re-expanded overnight with multiple washes in Milli-Q water.

Gels were mounted on 25 mm, no. 1.5H poly-L-lysine–coated coverslips in an imaging ring and imaged using a custom spinning disk microscope built on an Eclipse Ti-E body (Nikon) equipped with perfect focus, an MS-2000-XYZ stage with Piezo Top Plate (ASI), and a CSUX1-A1 spinning disk unit (Yokogawa). Imaging was performed with a 60×/1.2 NA water-immersion objective. Samples were excited using a 488 nm Voltran Stradus laser (150 mW), and emission was filtered with an ET525/50m (GFP) filter. Volume-scans of the expanded samples were acquired with a Prime BSI sCMOS camera (Teledyne Photometrics) at maximum laser intensity, using a 500 ms frame time and a z-step size of 200 nm. Microscope control was handled via MetaMorph 7.10 (Molecular Devices).

### Image and data analysis

#### Polarity Index of endogenous CSTN

We used a previously established method to investigate the polarized distribution of proteins, called the polarity index (PI) (Kapitein et al., 2010). Segmented lines were drawn along three dendrites using ImageJ and one portion of the axon of ∼30 μm (excluding the axon initial segment) in each image. The mean intensity of the three dendrites were subsequently averaged. Afterwards, the polarity index was measured using the following formula: PI = (Id − Ia)/(Id + Ia). Id indicates the mean intensity of the three dendrites and Ia indicates the mean intensity of the axon. PI <0 indicates axonal distribution, PI >0 indicates dendritic distribution and PI = 0 stands for non-polarized distribution where Id = Ia.

#### Kymograph analysis

Kymographs were generated from live-cell videos in ImageJ/FIJI. Segmented lines (30 μm) were drawn along axonal and dendritic tracks and subsequently straightened. Straightened axons and dendrites were then re-sliced followed by z-projection to obtain kymographs. Anterograde movements (towards tips) were oriented from left to right in all kymographs. Stationary, retrograde and anterograde compartments were manually counted, after which the percentage of compartments moving in each direction was calculated.

#### Quantification of the number of axonal compartments

To quantify vesicle number in the axon, we manually counted the number of puncta along a defined stretch of the axon (excluding the axon initial segment). RAB6A signal was additionally measured in the most distal tip of the axon. Intensity values were normalized to correct for differences in staining intensity between the replicates by multiplying each value by the overall mean intensity in the scrambled condition divided by the mean intensity in the scrambled condition of each respective replicate.

#### Colocalization analyses

To measure colocalization, we applied the ImageJ/FIJI plugin JACoP to obtain values for Pearson Correlation. Values were measured either in the soma or axons. Colocalization between cargoes in the axon was manually counted in a stretch of axon by counting the number of puncta containing individual and both markers.

#### Intensity measurements

To measure intensity (mean grey value) in the Golgi region, we generated a mask of the GM130 label using ImageJ/FIJI. Respective mask was subsequently used to measure the intensity of the marker of interest. To measure the secretion efficiency of RUSH cargoes, we analyzed the intensity of cargo signal in 40 µm segment of the axon (excluding the axon initial segment). Additionally, intensity in the perinuclear area was measured. The background intensity was subtracted from these measurements. Ratio was extracted by dividing intensity axon / intensity perinuclear area. The amount of RAB6A in the soma was quantified by drawing a polygon selection in the perinuclear area and measuring the mean intensity. This value was also normalized to correct for differences in staining intensity between the replicates by multiplying each value by the overall mean intensity in the scrambled condition divided by the mean intensity in the scrambled condition of each respective replicate.

#### Western blot analysis

Band intensities were measured in ImageJ/FIJI using the *Analyze > Gels* option. Extracted values were divided against loading control to correct for differences in total protein levels. Values are displayed as fold change.

#### Measurement of Golgi dispersal

Number of Golgi stacks was quantified by thresholding GM130 signal and analyzing the particle number using ImageJ. Particles smaller than 0.026 µm^2^ were removed from the analysis. Unspecific GM130 label in the nucleus was excluded from analyses.

#### Electron microscopy vesiculation

To assess the degree of vesiculation near the Golgi region in resin-embedded EM samples, a region of interest was selected at the trans side of the Golgi stack, matching the approximate size of the Golgi complex in the corresponding section. Endosomes as well as COPI- and COPII-coated vesicles were excluded from the analysis. Vesiculation was then scored according to the following scale: none (0%), low (0–10%), moderate (10–30%), and high (30–100%). Percentages represent the proportion of the area occupied by Golgi-derived vesicles, including both clathrin-coated and non-coated structures.

### Statistical analysis

Data processing and statistical analysis were performed using Microsoft Excel, GraphPad and Prism (version 9.5.1). Significance was determined as following: ns-not significant, *p < 0.05, **p < 0.01, ***p < 0.001. Data normality was checked using D’Agostino–Pearson omnibus test. The statistical test performed is indicated in figure legends and Source Data file. Exact p values, number of cells and number of replicates are reported in the Source Data file.

