## Supplementary Table 1 for "Calsyntenin-1 and calsyntenin-3 coordinate TGN exit of axonal cargoes"

**Table 1:** Primer table

| **Primer sequence** | **Cloning purpose** |
| --- | --- |
| cgtcagatccgctagcgctaccggtgccaccATGCTGCGCCGCCCCGCTC | pCMV_HaloTag-CSTN1 |
| cgatttccatgctaccGGCCCAGACCCCGCCGCC | pCMV_HaloTag-CSTN1 |
| ggccggtagcATGGAAATCGGTACTGGCTTTCCATTCGAC | pCMV_HaloTag-CSTN1 |
| taactcgcgcTGATCCGCCGCCACCCGA | pCMV_HaloTag-CSTN1 |
| cggcggatcaGCGCGAGTTAACAAGCAC | pCMV_HaloTag-CSTN1 |
| gcagaattcgaagcttgagctcgagTCAGTAGCTGAGGGTGGAGTC | pCMV_HaloTag-CSTN1 |
| cgtcagatccgctagcgctaccggtgccaccATGCTGCCTGGGCGGCTG | pCMV_HaloTag-CSTN2 |
| cgatttccatgctaccGCCGCTCCCCACGCCCAG | pCMV_HaloTag-CSTN2 |
| cggcggtagcATGGAAATCGGTACTGGCTTTCCATTCGAC | pCMV_HaloTag-CSTN2 |
| caccgccgctTGATCCGCCGCCACCCGA | pCMV_HaloTag-CSTN2 |
| cggcggatcaAGCGGCGGTGGCGGGGAC | pCMV_HaloTag-CSTN2 |
| gcagaattcgaagcttgagctcgagCTAGTAGGGGAGGGTGGAGTCATC | pCMV_HaloTag-CSTN2 |
| cgtcagatccgctagcgctaccggtgccaccATGACCCTCCTGCTGCTG | pCMV_HaloTag-CSTN3 |
| cgatttccatgctaccACAGGAGCAGGACGC | pCMV_HaloTag-CSTN3 |
| gctcctgtggtagcATGGAAATCGGTACTGGCTTTCCATTCGAC | pCMV_HaloTag-CSTN3 |
| tggctttgttTGATCCGCCGCCACCCGA | pCMV_HaloTag-CSTN3 |
| cggcggatcaAACAAAGCCAACAAGCACAAG | pCMV_HaloTag-CSTN3 |
| gcagaattcgaagcttgagctcgagTTAGTAGCGGTGTGGGGG | pCMV_HaloTag-CSTN3 |
| tgctcaccatgctaccGGCCCAGACCCCGCCGCC | pCMV_mNeonGreen-CSTN1 |
| ggccggtagcATGGTGAGCAAGGGCGAG | pCMV_mNeonGreen-CSTN1 |
| cgccaccagaCTTGTACAGCTCGTCCATGC | pCMV_mNeonGreen-CSTN1 |
| gctgtacaagTCTGGTGGCGGAGGCTCG | pCMV_mNeonGreen-CSTN1 |
| gcagaattcgaagcttgagctcgagTCAGTAGCTGAGGGTGGAGTCATC | pCMV_mNeonGreen-CSTN1 |
| tgctcaccatgctaccACAGGAGCAGGACGC | pCMV_mNeonGreen-CSTN3 |
| GCTCCTGtggtagcATGGTGAGCAAGGGCGAG | pCMV_mNeonGreen-CSTN3 |
| gagagctgagtcatccgcgaagaggtctgggtcc | pCMV_mNeonGreen-CSTN3_KBS |
| ggacccagacctcttcgcggatgactcagctctc | pCMV_mNeonGreen-CSTN3_KBS |
| ATCGAAAATACCGAAAAA CTGCAGTACAGTGGTGAGAGG | pCMV_mNeonGreen-CSTN3_shResistant |
| TTTTTCGGTATTTTCGATGTTCCCGTCATTGTCAATGAGG | pCMV_mNeonGreen-CSTN3_shResistant |
| ttgaaaaacacgatgataagaattcaaATGTACAGGATGCAACTC | FUGW_Strep-KDEL_IL2-SBP-TRKB-mNG |
| acatggggcagctgccgctgccgctgccTGGTTCACGTTGACCTTG | FUGW_Strep-KDEL_IL2-SBP-TRKB-mNG |
| acgtgaaccaggcagcggcagcggcagcTGCCCCATGTCCTGCAAATg | FUGW_Strep-KDEL_IL2-SBP-TRKB-mNG |
| tgccgctaccCCCTAGGATGTCCAGGTAGAC | FUGW_Strep-KDEL_IL2-SBP-TRKB-mNG |
| catcctagggGGTAGCGGCAGCGGTAGCatg | FUGW_Strep-KDEL_IL2-SBP-TRKB-mNG |
| ttgatatcgaattgttaacggatccTTACTTGTACAGCTCGTCCATGCC | FUGW_Strep-KDEL_IL2-SBP-TRKB-mNG |
| AAAACACGATGATAAgAATTATGGTCGTGATGCTGCGGTA | FUGW_Strep-Ii_L1CAM-mNG-SBP |
| GCAGCaGAtCCAGCGGATCCTTCTAGGGCTACTGCAGGATTG | FUGW_Strep-Ii_L1CAM-mNG-SBP |
| GGATCCGCTGGaTCtGCTGCAGGTTCTGGCGCTGGcTCcGCTGCTGGTTCT | FUGW_Strep-Ii_L1CAM-mNG-SBP |
| CTGGcTCcGCTGCTGGTTCTGGCGAATTCGTGAGCAAGGGCGAGGAGGA | FUGW_Strep-Ii_L1CAM-mNG-SBP |
| CTTGTACAGCTCGTCCATGCC | FUGW_Strep-Ii_L1CAM-mNG-SBP |
| GGCATGGACGAGCTGTACAAGGGATCCGACGAGAAGACCACTGGTTGG | FUGW_Strep-Ii_L1CAM-mNG-SBP |
| GAATTGTTAACGGATCGAATTCTTATGGTTCACGTTGACCTTGTGG | FUGW_Strep-Ii_L1CAM-mNG-SBP |
| ttgaaaaacacgatgataaggccaccATGTACAGGATGCAACTCCTGTC | pFSW-StrepKDEL_SBP-TRKB-mNG |
| ttgaaaaacacgatgataaggccaccATGCTGCCCGGTTTGGCAC | pFSW-StrepKDEL_SBP-TRKB-mNG |
| tatcgaattgttaacggatcTTAGTTCTGCATCTGCTCAAAGAACTTG | pFSW-Strep-KDEL_SBP-mNG-APP |
| TATCGAATTGTTAACGGATCTTACTTGTACAGCTCGTCCATGCC | pFSW-Strep-KDEL_SBP-mNG-APP |
| gcttgggctgcaggtcgactgccaccatgCGGATCCGGGCCGCACAT | FUGW_CtermCSTN1-GFP |
| tgccgctaccGTAGCTGAGGGTGGAGTCATCCC | FUGW_CtermCSTN1-GFP |
| cctcagctacGGTAGCGGCAGCGGTAGC | FUGW_CtermCSTN1-GFP |
| tcgaattgttaacggatccgTTACTTGTACAGCTCGTCCATGCC | FUGW_CtermCSTN1-GFP |
| gcttgggctgcaggtcgactgccaccatgCGCATCCATTCCCTTCACCG | FUGW_CtermCSTN3-GFP |
| tgccgctaccGTAGCGGTGTGGGGGGGT | FUGW_CtermCSTN3-GFP |
| acaccgctacGGTAGCGGCAGCGGTAGC | FUGW_CtermCSTN3-GFP |
| AAAACACGATGATAAgAATTatgtaccagaggatgctccg | FUGW_Strep-Ii_NRXN1B-mNG-SBP |
| GTGGTCTTCTCGTCACCGGTcccccaggccactcctagg | FUGW_Strep-Ii_NRXN1B-mNG-SBP |
| CTGCTGGTTCTGGCGAATTCGCATCCAGTTTGGGAGCGCA | FUGW_Strep-Ii_NRXN1B-mNG-SBP |
| GCAGCGGAGCCAGCGGATCCGACATAATACTCCTTATCCTTGTTCTTCTTG | FUGW_Strep-Ii_NRXN1B-mNG-SBP |
| GGATCCGCTGGCTCCGCTGCTGGTTCTGGCGAATTCGACGAGAAGACCACTGGTTG | FUGW_Strep-Ii_NRXN1B-mNG-SBP |
| GGGTTAGGGATAGGCTTACCGGATCCTGGTTCACGTTGACCTTGTG | FUGW_Strep-Ii_NRXN1B-mNG-SBP |
| GTGAGCAAGGGCGAGGAGGA | FUGW_Strep-Ii_NRXN1B-mNG-SBP |
| TCGAATTGTTAACGGATCGAATTCTTACTTGTACAGCTCGTCCATGCC | FUGW_Strep-Ii_NRXN1B-mNG-SBP |
| ttgggctgcaggtcgactctagagccaccatgcacaggaggagaagcag | FUGW_Strep-Ii_L1CAM-mNG-SBP |
| gctagcttcgaagaattcttatcatcgtgtttttcaaaggaaaacca | FUGW_Strep-Ii_L1CAM-mNG-SBP |
| CAACTCCCTCAAGATTGTCAGCAA | qPCR GAPDH |
| GGCATGGACTGTGGTCATGA | qPCR GAPDH |
| GTTCTGGGAAACTGGCAGAT | qPCR CSTN1 |
| TCTGTTGCCATCCTCAGGG | qPCR CSTN1 |
| GACCTCTTGCCATCCCCTAG | qPCR CSTN2 |
| CCTGCCTGCCATCAAACTTG | qPCR CSTN2 |
| AGTGCTGCGACTCTCATCAT | qPCR CSTN3 |
| GGCGATGAAGGGAGTGGAT | qPCR CSTN3 |
| tgtgACACTCCTAAGGAATCAGC | CSTN1 CRISPIE_1 guide |
| aacGCTGATTCCTTAGGAGTGTc | CSTN1 CRISPIE_1 guide |
| gcgACACTCCTAAGGAATCAGCggg | CSTN1 CRISPIE_1 5'donor |
| agccccGCTGATTCCTTAGGAGTGT | CSTN1 CRISPIE_1 5'donor |
| ggaACACTCCTAAGGAATCAGCggg | CSTN1 CRISPIE_1 3'donor |
| cgtcccGCTGATTCCTTAGGAGTGT | CSTN1 CRISPIE_1 3'donor |
| tgtgGCGACGGCGGGGACAGCCGG | CSTN2 ORANGE2 guide |
| aacCCGGCTGTCCCCGCCGTCGCc | CSTN2 ORANGE2 guide |
| gcgcccCCGGCTGTCCCCGCCGTCGCt | CSTN2 ORANGE2 5'donor |
| agcaGCGACGGCGGGGACAGCCGGggg | CSTN2 ORANGE2 5'donor |
| ggacccCCGGCTGTCCCCGCCGTCGC | CSTN2 ORANGE2 3'donor |
| cgtGCGACGGCGGGGACAGCCGGggg | CSTN2 ORANGE2 3'donor |
| tgtgGAGCTAGGAGGACCATCCTG | CSTN3 CRISPIE_1 guide |
| aacCAGGATGGTCCTCCTAGCTCc | CSTN3 CRISPIE_1 guide |
| gcgGAGCTAGGAGGACCATCCTGggg | CSTN3 CRISPIE_1 5'donor |
| agccccCAGGATGGTCCTCCTAGCTC | CSTN3 CRISPIE_1 5'donor |
| ggaGAGCTAGGAGGACCATCCTGggg | CSTN3 CRISPIE_1 3'donor |
| cgtcccCAGGATGGTCCTCCTAGCTC | CSTN3 CRISPIE_1 3'donor |
