## Supplementary Table 2 for "Calsyntenin-1 and calsyntenin-3 coordinate TGN exit of axonal cargoes"

**Table 2:** Antibody table

| **Name** | **Host** | **Catalog number** | **Company** | **Dilution** |
| --- | --- | --- | --- | --- |
| Primary antibodies | | | | |
| APP (C. term) | Rabbit | ab32136 | Abcam | 1:200 (IF) |
| GFP | Mouse | G1546 | Sigma | 1:2000 (WB) |
| RFP / mCherry | Rabbit | 600-401-379 | Rockland | 1:2000 (WB) |
| CSTN3 | Rabbit | 13302-1-AP | Proteintech | 1:1000 (WB) |
| GM130 | Mouse | 610823 | BD Biosciences | 1:500 (IF) |
| GM130 | Rabbit | ab52649 | Abcam | 1:800 (IF) |
| RAB6A | Mouse | Clone 5B10 | Gift from Angelika Barnekow | 1:100 (IF)  1:500 (WB) |
| TGN38 | Rabbit | ab16059 | Abcam | 1:2000 (IF) |
| TRIM46 | Rabbit |  | In house | 1:1000 (IF) |
| α-tubulin | Mouse | T5168 | Sigma | 1:20000 (WB) |
| Actin | Mouse | 0869100-CF | MP Biochemicals | 1:20000 (WB) |
| ALFA nanobody | Rabbit | N1583 | NanoTag Biotechnologies | 1:500 |
| Secondary antibodies | | | | |
| Anti-mouse HRP | Rabbit | P0260 | Agilent | 1:10000 (WB) |
| Anti-rabbit HRP | Swine | P0399 | Agilent | 1:10000 (WB) |
| anti-mouse IRDye 680 | Donkey | 926-68,020 | LICORbio | 1:20000 (WB) |
| Anti-mouse Alexa 405 | Goat | A31553 | Life Technologies | 1:500 (IF) |
| Anti-rabbit Alexa 405 | Goat | A31556 | Life Technologies | 1:500 (IF) |
| Anti-Mouse Alexa 488 | Goat | A11029 | Life Technologies | 1:1000 (IF) |
| Anti-Rabbit Alexa 488 | Goat | A11034 | Life Technologies | 1:1000 (IF) |
| Anti-Mouse Alexa 647 | Goat | A21236 | Life Technologies | 1:1000 (IF) |
| Anti-Rabbit Alexa 647 | Goat | A21245 | Life Technologies | 1:1000 (IF) |
